# Sensory lesioning induces microglial synapse elimination via ADAM10 and fractalkine signaling

**DOI:** 10.1101/551697

**Authors:** Georgia Gunner, Lucas Cheadle, Kasey M. Johnson, Pinar Ayata, Ana Badimon, Erica Mondo, Aurel Nagy, Liwang Liu, Shane M. Bemiller, Ki-Wook Kim, Sergio A. Lira, Bruce T. Lamb, Andrew R. Tapper, Richard M. Ransohoff, Michael E. Greenberg, Anne Schaefer, Dorothy P. Schafer

**Author notes:** Corresponding Author: Dorothy (Dori) Schafer. Authors contributed equally (Lucas Cheadle, Kasey M. Johnson).

## Abstract

Microglia rapidly respond to changes in neural activity and inflammation to regulate synaptic connectivity. The extracellular signals, particularly neuron-derived molecules, that drive these microglial functions at synapses remains a key open question. Here, whisker lesioning, known to dampen cortical activity, induces microglia-mediated synapse elimination. We show that this synapse elimination is dependent on the microglial fractalkine receptor, CX3CR1, but not complement receptor 3, signaling. Further, mice deficient in the CX3CR1 ligand (CX3CL1) also have profound defects in synapse elimination. Single-cell RNAseq then revealed that *Cx3cl1* is cortical neuron-derived and ADAM10, a metalloprotease that cleaves CX3CL1 into a secreted form, is upregulated specifically in layer IV neurons and microglia following whisker lesioning. Finally, inhibition of ADAM10 phenocopies *Cx3cr1*^-/-^ and *Cx3cl1*^-/-^ synapse elimination defects. Together, these results identify novel neuron-to-microglia signaling necessary for cortical synaptic remodeling and reveal context-dependent immune mechanisms are utilized to remodel synapses in the mammalian brain.

## INTRODUCTION

Microglia are constant surveyors of their extracellular environment and dynamic regulators of synaptic connectivity. Among the recently identified functions of microglia at synapses is developmental synaptic pruning, whereby microglia are, in fact, ‘listening’ to circuit activity and engulfing synapses from less active neurons^1,2^. There has also been recent work in the context of the inflamed central nervous system (CNS) where microglia engulf and eliminate synapses during neurodegeneration^3-5^, suggesting that similar developmental mechanisms are upregulated at the wrong time and place during disease. Mechanisms regulating this process of microglia-mediated synapse elimination in health and disease have largely focused on surface receptors expressed by microglia. Whether there are neuronal cues that instruct microglia to eliminate synapses remains an open question. Given the large array of neurological disorders that implicate microglia-mediated synapse elimination as a regulator of synaptic connectivity, understanding the underlying neuron-microglia signaling will be critical going forward.

Two of the major molecular pathways identified to modulate microglia function at synapses are phagocytic signaling through complement receptor 3 (CR3) and chemokine signaling through the fractalkine receptor (CX3CR1). In the developing mouse visual thalamus, complement proteins C3 and C1q localize to synapses and microglia engulf synapses via CR3 expressed by microglia^1,6,7^. Blocking this synaptic engulfment in C3, C1q, or CR3-deficient mice results in sustained synaptic pruning defects. Similar complement-dependent mechanisms of synapse elimination have been identified in mouse models of neurodegeneration^3-5^. CX3CR1 is a G-protein coupled chemokine receptor highly enriched in microglia^8^. While CR3-dependent phagocytic signaling regulates synaptic pruning in the developing visual system, studies have demonstrated that these effects are independent of CX3CR1^9,10^. Instead, in the developing hippocampus and barrel cortex, CX3CR1-deficient mice exhibit a transient delay in microglial recruitment to synapse-dense brain regions and a concomitant delay in functional maturation of synapses^11,12^. Long term, CX3CR1-deficient mice demonstrate defects in social interactions and functional synaptic connectivity^13^. How CX3CR1 deficiency is exerting these effects and the relative involvement of its canonical ligand fractalkine (CX3CL1) are unknown.

In the current study, we used the mouse barrel cortex system to identify mechanisms by which neurons communicate with microglia to regulate synapse remodeling. Sensory endings from trigeminal neurons innervate the whisker follicles on the snout. Sensory information from each whisker is then transmitted to the brain stem, then to the ventral posteromedial (VPM) nucleus of the thalamus, and ultimately to the cortex via thalamocortical (TC) synapses, which terminate largely in layer IV of the barrel cortex. This is a particularly powerful system for studying synapse remodeling as each whisker corresponds to an individual bundle of TC inputs (i.e. barrels) separated by septa, which, despite being several synapses away, are highly sensitive to manipulation of the whiskers^14,15^. Indeed, removal of the whiskers in this system results in dampened activity in the cortex and elimination of TC synapses^16-28^. However, the mechanism(s) by which changes in activity elicit TC synapse remodeling is in an open question.

To address these open questions, we used whisker cauterization and trimming in developing mice, paradigms known to reduce activity in the corresponding barrel cortex^16-28^. We identify synapse elimination within 1 week of whisker removal and robust microglia-mediated synaptic engulfment. Unlike the developing visual system, synapse elimination in the barrel cortex is CR3-independent. Instead, we identify profound defects in TC synapse elimination in mice deficient in CX3CR1 enriched in microglia or its ligand CX3CL1. Using single-cell sequencing, we further uncover that CX3CL1 is enriched in cortical neurons, but its transcription is not modulated by sensory lesioning. Instead, ADAM10, a metalloprotease known to cleave CX3CL1 into a secreted form is increased in layer IV neurons and microglia. Strikingly, pharmacological inhibition of ADAM10 phenocopies defects in sensory lesion-induced TC synapse remodeling in CX3CL1 and CX3CR1-deficient mice. This suggests that post-translational modification of neuronal CX3CL1 by ADAM10 following loss of sensory input directs microglia to eliminate synapses. Together, these data provide new insight into molecular mechanisms by which neurons communicate with microglia and direct them to engulf and remove synapses. Our single-cell RNAseq further provides an unbiased approach to identify novel mechanisms regulating neuron-microglia communication in response to sensory perturbations. We also demonstrate that distinct immune signaling pathways are engaged by microglia to remodel synapses in different contexts and, in the process, uncover a new molecular pathway regulating remodeling of TC synapses.

## RESULTS

### Whisker lesioning induces rapid and robust elimination of thalamocortical inputs in the barrel cortex

To interrogate neuron-microglia signaling regulating synapse remodeling, we performed two different manipulations in separate cohorts of postnatal day 4 (P4) mice, unilateral whisker trimming and unilateral whisker lesioning by cauterization (P4; Fig. 1 and Supplemental Fig.1a-b). These paradigms are known to reduce activity in the barrel cortex and maximize robust TC remodeling in the neonate while avoiding the critical window (P0-P3) where sensory loss disrupts initial TC input wiring and can induce neuronal apoptosis in the barrel cortex^16-18,28^. Also, each model has its own internal control as the whiskers are left intact on the other side of the snout. Using anti-vesicular glutamate transporter 2 (VGluT2) immunostaining, we observed a decrease in TC presynaptic terminals within layer IV of the barrel cortex by 17 days-post the beginning of the whisker trimming paradigm (Supplemental Fig.1a-b). These results are consistent with previous work showing decreased TC inputs within the barrel cortex following whisker trimming in neonates and adults^21,22,27^. This decrease in presynaptic terminals was accelerated in the whisker lesioning model where, within 6 days, VGluT2 immunoreactivity was decreased by ∼75% in the deprived compared to the control barrel cortex within the same animal (Fig. 1b-c). Demonstrating that these results were not due to a downregulation of VGluT2, we observed similar effects in mice that express a fluorescent reporter in thalamic neurons and their cortical projections (*SERT-Cre; Rosa26*^*LSL.TdTomato/+*^) (Supplementary Fig. 1c-d). To note, presynaptic terminals and axons are labeled with tdTomato as well as other cortical projections besides TC synapses in SERT-Cre;*Rosa26*^*LSL.TdTomato/+*^ mice, resulting in less robust detection of synapse elimination compared to analyses with VGluT2, a more specific marker for TC presynaptic terminals.

**Figure 1.**
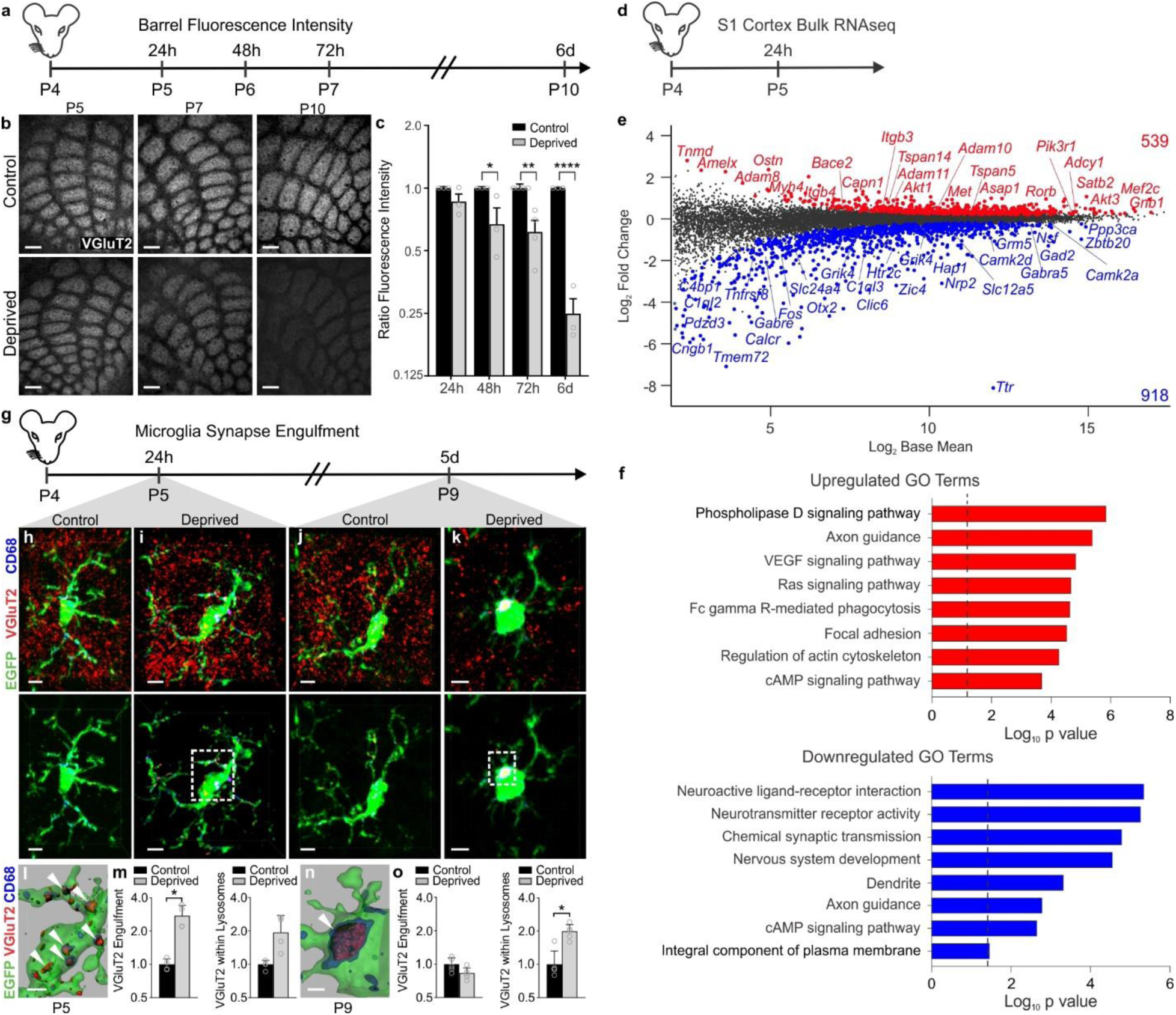
Whisker lesioning induces microglial engulfment and elimination of TC inputs within the barrel cortex. **a**, Timeline for analysis of TC input elimination following whisker lesioning at P4. **b**, Tangential sections of layer IV contralateral control (top panel) and deprived (bottom panel) barrel cortices immunolabeled for anti-VGluT2 show a decrease in TC inputs by P10. Scale bar, 150 µm. **c**, Quantification of fluorescence intensity of VGluT2-positive TC inputs in the barrel cortex in the deprived (gray bars) compared to the control barrel cortex (black bars) at each time point post-whisker removal. Data normalized to the control, non-deprived hemisphere within each animal. (Two-way ANOVA with Sidak’s post hoc; control vs deprived 24h, n = 3 animals, *P* = 0.5323, *t* = 1.419, *df* = 18; control vs deprived 48h, n = 3 animals, *P* = 0.0142, *t* = 3.349, *df* = 18; control vs deprived 72h, n = 4 animals, *P* = 0.0011, *t =* 4.516, *df* = 18; control vs deprived 6d, n = 3 animals, *P* <0.0001, *t =* 7.631, *df* = 18). **d,** Timeline for bulk RNAseq of the barrel cortex 24 hours after whisker lesioning**. e,** Quantification for the Log_2_ Fold Change for genes enriched in the deprived somatosensory cortex. Upregulated genes annotated in red, downregulated genes annotated in blue. **f**, Gene ontology analysis across genes significantly up (red) or down (blue) regulated in the barrel cortex 24 hours after whisker lesioning. **g,** Timeline for analysis of TC input engulfment by microglia. **h,i,j,k**, Fluorescent images of microglia (green) within layer IV of the control (h,j) and deprived (i,k) barrel cortices 24 h (h,i) and 5 d (j,k) after unilateral whisker removal. Microglial lysosomes are labeled with anti-CD68 (blue). Raw fluorescent images, top panel; VGlut2 signal internalized within microglia, bottom panel. Scale bar, 5 µm. **l,n**, 3D surface-rendered inset of i,k (bottom panel). Arrows depict VGlut2 (red) internalized within microglia (green) and within lysosomes (blue). Scale bar, 2 µm. **m,o**, Quantification of VGlut2 engulfment within microglia (left) and VGlut2 engulfment within lysosomes (right) 24 h (m; within microglia, Two-sided Student’s t-test, n = 4 animals, *P* = 0.0305, *t* = 2.642 *df* =6; within lysosomes, Two-sided Student’s t-test, n = 4 animals, *P* = 0.2955, *t* = 1.146, *df* =6) and 5 d (o; within microglia, Two-sided Student’s t-test, n = 5 animals, *P* = 0.3319, *t* =1.033, *df* =8; within lysosomes, Two-sided Student’s t-test, n = 5 animals, *P* = 0.0272, *t* =2.251, *df* =8) after whisker removal in *Cx3cr1*^*EGFP/+*^ microglia reveals increased VGluT2 within microglia at 24 h (m) and increased VGluT2 within microglial lysosomes at 5 d-post whisker removal (o). Data normalized to engulfment in microglia in the control hemisphere within each animal. All data presented as mean ± SEM.

To determine if TC synapse elimination induced by whisker lesioning was an indirect effect, due to neuronal damage and loss, we also assessed cell death, axon degeneration, and cell stress. We observed no significant increase in cleaved Caspase3+ neurons or amyloid precursor protein (APP) accumulation in axons within the barrel cortex circuit following whisker removal (Supplementary Figure 2d-f). These analyses included neurons within the VPM nucleus of the thalamus, cortical neurons within the barrel cortex, and trigeminal neurons, which innervate the whisker follicle. However, an increase in the stress marker ATF3 was observed in trigeminal neurons (Supplementary Fig. 2a-b), but not in cortical or VPM neurons (Supplementary Fig. 2c). To further characterize the system, we performed bulk RNAseq of the barrel cortex following whisker lesioning. Consistent with decreased activity in the barrel cortex, immediate early genes such as *Fos* and genes related to neurotransmitter signaling were decreased in the deprived barrel cortex with no apparent increase in genes related to cell stress or death (Fig. 1d-f; Fig. 8j-k). Instead, there was an increase in genes related to axon growth and phagocytic signaling. These data are most consistent with synapse loss resulting from decreased activity in the barrel cortex circuit vs. injury-induced inflammation and degeneration. Given the rapid and robust presynaptic terminal loss elicited in the absence of significant neuronal cell death or degeneration, we proceeded with whisker cauterization-induced sensory lesioning in P4 mice as a model for the remainder of the study.

### Whisker lesioning induces microglial engulfment of TC inputs within the barrel cortex independent of complement receptor 3 (CR3)

In the developing visual system, microglia engulf and remove synapses in response to dampened neural activity^1,2,29^. Therefore, we next explored whether microglia similarly engulf and eliminate TC synaptic inputs in the barrel cortex following whisker lesioning. Microglia were labeled using a transgenic mouse that expresses EGFP under the control of the fractalkine receptor (*Cx3cr1*^*EGFP/+*^)^30^ and TC presynaptic terminals were labeled using an anti-VGluT2 antibody. Using fluorescent confocal microscopy and structured illumination microscopy, we detected an ∼2-fold increase in the volume of engulfed TC inputs within microglia in the deprived barrel cortex within 24 h of removing the whiskers (Fig. 1g-o and Supplementary Fig. 3a)^1,31^. These inputs were largely localized within the microglia, but not yet significantly associated with lysosomes (Fig. 1g-i). Interestingly, at 5 days-post whisker removal, microglia had a more phagocytic morphology, with a more rounded and enlarged soma, and most engulfed TC inputs were completely localized within microglial lysosomes in the deprived barrel cortex (Fig. 1k-o). In contrast, minimal TC inputs were detected within microglia or microglial lysosomes in the control barrel cortex of the same animal at any time point assessed (Fig. 1h,j). In addition, we observed, not only an increase in the amount of synaptic material engulfed, but the percentage of highly phagocytic microglia (phagocytic index >1% at 24 h and >2% at 5d) was also increased in the deprived cortex (Supplementary Fig. 3). This increase in phagocytic activity was further reflected in our RNAseq data (Fig. 1e-f). These data provide the first evidence that microglia engulf and eliminate TC inputs in the cortex several synapses away from a peripheral sensory lesion known to reduce activity in the cortex.

One of the best characterized mechanisms utilized by microglia to engulf and remodel synapses is complement-dependent phagocytic signaling^1,6,7^. During synaptic pruning in the developing mouse visual system, complement proteins C1q and C3 localize to synapses. Microglia subsequently engulf and eliminate these synapses via the microglial phagocytic receptor complement receptor 3 (CR3/CD11b). To determine whether this pathway also regulates microglia-mediated synapse elimination in the barrel cortex following whisker lesioning, we assessed microglial engulfment and elimination of TC inputs in CR3-deficient (CR3-KO) mice. Surprisingly, TC inputs were eliminated appropriately by 6 days-post whisker removal in CR3-KO mice (Fig. 2a-c) and engulfed TC inputs were detected within microglia in the deprived barrel cortex (Fig. 2d-f), as described above for wild-type mice (Fig. 1). Together, these data demonstrate that CR3 is dispensable for sensory lesion-induced microglial engulfment and elimination of TC synapses in the barrel cortex.

**Figure 2.**
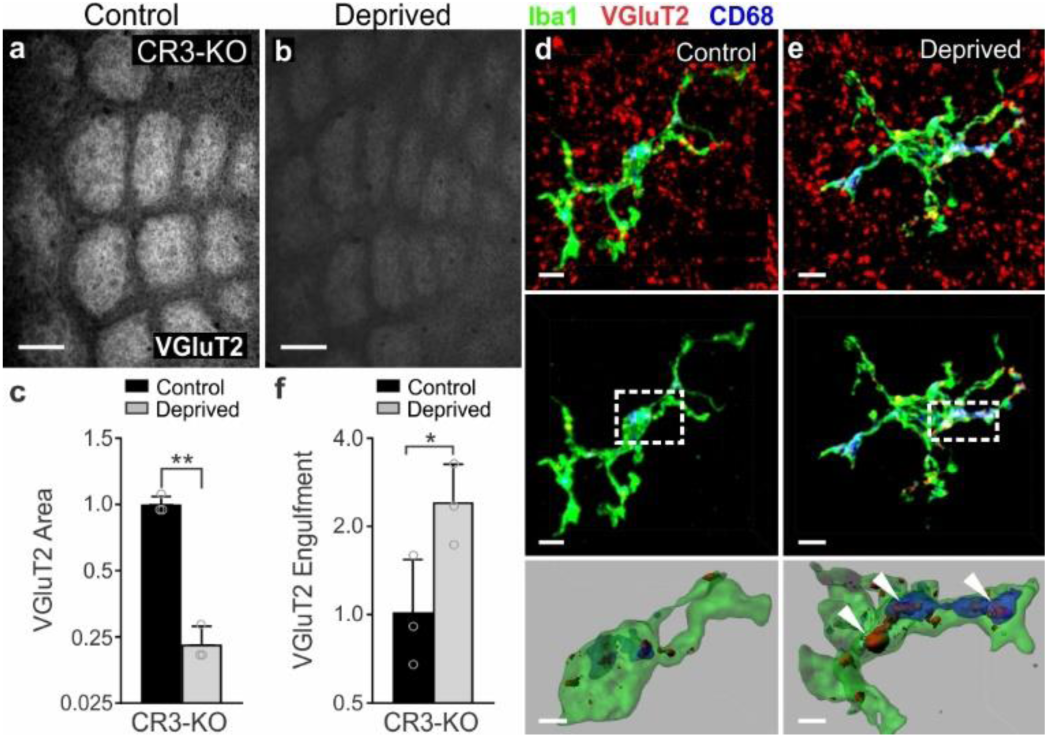
CR3 is not required for whisker lesion-induced TC input engulfment and elimination. **a,b**, Representative images of VGluT2 immunoreactivity in the non-deprived (a) and deprived (b) barrel cortices of CR3-KO mice show TC inputs are properly eliminated 6 d after whisker lesioning. Scale bar, 150 µm. **c**, Quantification of VGluT2 area within barrel centers 7 d after whisker removal reveals that TC inputs are still eliminated in CR3-KO mice. (Data normalized to the control, non-deprived VGluT2 area within each animal; Two-tailed Student’s t-test, n = 3 animals, *P* = 0.0015, *t* =7.765 *df* =4). **d,e**, Top Panels, Fluorescent images of microglia within the control (d) and deprived (e) barrels of CR3-KO mice. Middle panels depict VGluT2 signal (red) within microglia (green) and within lysosomes (blue). Bottom panels are 3D surface-rendered insets (dotted boxes in middle panels) of control (d) and deprived (e) CR3-KO microglia. Arrows depict increased VGluT2 internalization within microglia in the deprived hemisphere. Scale bar, 2 µm. Scale bar, 5 µm. **f**, Quantification of VGluT2 engulfment within CR3-KO microglia 24 h after whisker removal reveals that CR3 deficiency fails to block microglial engulfment of TC inputs. (Data normalized to engulfment in microglia in the control hemisphere within each animal; Two-tailed Student’s t-test, n = 3 animals, *P =* 0.260, *t* =4.215 *df* =2). All data presented as mean ± SEM.

### CX3CR1-deficient mice have profound defects in structural and functional synapse remodeling following whisker lesioning

Because of the surprising result that microglial engulfment and elimination of TC inputs in the barrel cortex was CR3-independent, we next explored other candidate mechanisms. Besides CR3, the fractalkine receptor (CX3CR1), a G-protein coupled chemokine receptor highly enriched in microglia, has also been implicated in regulating microglial function at developing synapses^11-13^. We, therefore, sought to assess whisker lesion-induced TC synapse remodeling in *Cx3cr1*^-/-^ mice. Remarkably, unlike CR3-KO mice (Fig. 2), elimination of VGluT2-positive TC presynaptic inputs and structural synapses (co-localized presynaptic VGluT2 and postsynaptic Homer puncta) 6 days-post whisker removal was completely blocked in *Cx3cr1*^-/-^ mice compared to *Cx3cr1*^*+*|*-*^ or *Cx3cr1*^*+*|*+*^ littermates (Fig. 3). This was largely a presynaptic effect as the density of postsynaptic Homer-positive immunoreactivity was relatively unaffected 6 days-post whisker removal, even in wild-type animals (Fig. 3h-i,l). This is consistent with previous work demonstrating that microglia preferentially engulf presynaptic inputs^1,32^. We further measured macrophage recruitment to the whisker follicles following whisker lesioning as well as immunostaining for ATF3 in *Cx3cr1*^-/-^ mice (Supplementary Fig. 4). All these responses were comparable to *Cx3cr1*^*+*|*-*^ controls (Supplementary Fig. 2). Last, we assessed TC input elimination following whisker trimming and found that TC input elimination elicited in this paradigm was also CX3CR1-dependent (Supplementary Fig. 4f-g). Together, these results suggest that synapse elimination defects in *Cx3cr1*^-/-^ mice were not secondary to changes in injury, wound healing, or neuronal stress responses.

**Figure 3.**
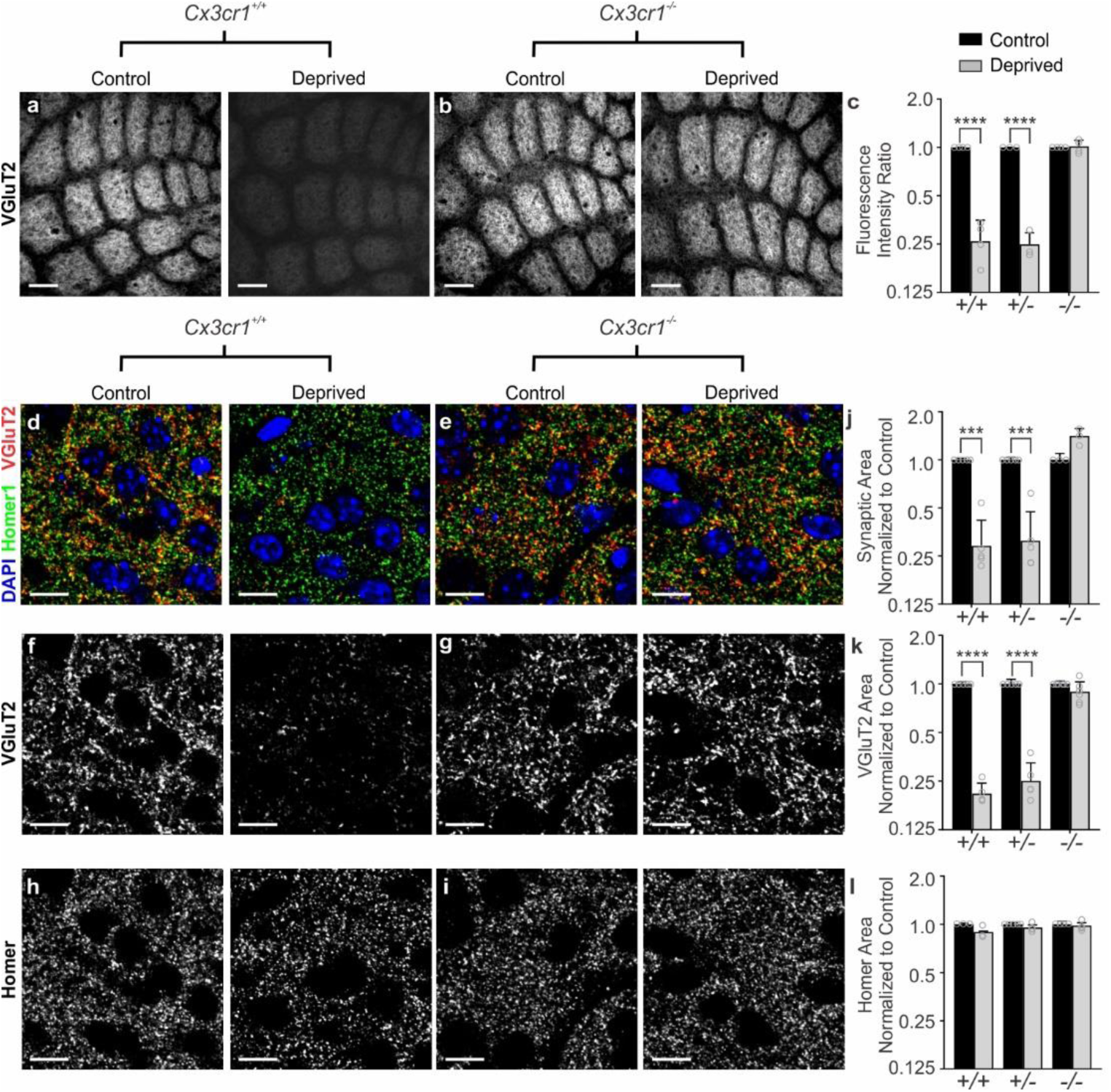
*Cx3cr1* expression is necessary for TC input elimination after whisker lesioning. **a-b,** VGluT2 immunolabeled TC inputs within tangential sections of control and deprived barrel cortices in *Cx3cr1*^+/+^ (a) and *Cx3cr1*^-/-^ mice (b) show TC inputs remain 6 d-post deprivation in mice lacking *Cx3cr1*. Scale bar, 150 µm**. c**, Quantification of fluorescence intensity of VGluT2-positive TC input immunoreactivity 6 d after deprivation in *Cx3cr1*^+/+^, *Cx3cr1*^+/-^ and *Cx3cr1*^-/-^ littermates demonstrates a significant decrease in VGluT2 immunoreactivity in *Cx3cr1*^+/+^, *Cx3cr1*^+/-^ mice following deprivation but this is blocked in *Cx3cr1*^-/-^ littermates. (Data normalized to the control, non-deprived hemisphere within each animal; Two-Way ANOVA with Sidak’s post hoc; control vs deprived *Cx3cr1*^+/+^ n = 4 animals, *P* < 0.0001, *t* = 8.967, *df* =16; control vs deprived *Cx3cr1*^+/-^ n = 3 animals, *P* < 0.0001, *t =* 7.882, *df* = 16; control vs deprived *Cx3cr1*^-/-^ n = 4 animals, *P* = 0.9976, *t* = 0.1722, *df* = 16). **d-i**, High magnification (63X) confocal images of TC synapses within layer IV of the control and deprived barrel cortices immunolabeled with presynaptic anti-VGluT2 (red) and postsynaptic anti-Homer (green) 6 d-post deprivation in *Cx3cr1*^+/+^ (d,f,h) and *Cx3cr1*^-/-^ (e,g,i) mice. Merged channels are shown in panels d-e. The presynaptic VGluT2 channel alone is shown in panels f-g. The postsynaptic Homer channel alone is shown in panels h-i. Scale bars, 10 µm. **j-l**, Quantification of d-i reveals a significant decrease in structural synapses (j; colocalized VGluT2 and Homer, Two-Way ANOVA with Sidak’s post hoc, control vs deprived *Cx3cr1*^+/+^ n = 5 animals, *P* = 0.0004, *t* = 4.765, *df* =20; control vs deprived *Cx3cr1*^+/-^ n = 5 animals, *P* = 0.0005, *t =* 4.617, *df* = 20; control vs deprived *Cx3cr1*^-/-^ n = 3 animals, *P* = 0.1290, *t* = 2.139, *df* = 20) and VGluT2-positive TC presynaptic terminal density (k; VGlut2 Area, Two-Way ANOVA with Sidak’s post hoc, control vs deprived *Cx3cr1*^+/+^ n = 5 animals, *P* < 0.0001, *t* = 6.919, *df* =26; control vs deprived *Cx3cr1*^+/-^ n = 5 animals, *P* <0.0001, *t =* 6.552, *df* = 26; control vs deprived *Cx3cr1*^-/-^ n = 6 animals, *P* = 0.6907, *t* = 1.006, *df* = 26) in *Cx3cr1*^+/+^ and *Cx3cr1*^+/-^ mice, which was blocked in *Cx3cr1*^-/-^ littermates. There was no significant change in postsynaptic Homer density (l; Homer Area, Two-way ANOVA with Sidak’s post hoc, control vs deprived *Cx3cr1*^+/+^ n = 3 animals, *P* = 0.2386, *t* = 1.811, *df* =18; control vs deprived *Cx3cr1*^+/-^ n = 5 animals, *P* =0.7852, *t =* 0.9918, *df* = 18; control vs deprived *Cx3cr1*^-/-^ n = 4 animals, *P* = 0.9731, *t* = 0.3908, *df* = 18) in any genotype. Data normalized to the control, non-deprived cortex within each animal. All data presented as mean ± SEM.

To further determine if CX3CR1-dependent defects in TC input elimination are long-lasting, we lesioned whiskers at P4 and then assessed structural and functional remodeling at ≥6 weeks of age (Fig. 4). Similar to 6 days post-whisker removal, we observed a significant decrease in structural TC inputs in the deprived adult (P90) *Cx3cr1*^+/+^ barrel cortex (Fig. 4a), which was blocked in *Cx3cr1*^-/-^ mice (Fig.4b). To determine whether structural synapse remodeling defects translated to changes in functional connectivity, we next performed electrophysiological recordings in ≥6 week *Cx3cr1*^-/-^ and *Cx3cr1*^*+*l^*+* mice. We observed a significant decrease in spontaneous excitatory postsynaptic current (sEPSC) frequency and amplitude in layer IV stellate neurons within the deprived cortex in *Cx3cr1*^+/+^ mice (Fig. 4c-d), which is consistent with a decrease in the number and strength of functional synapses in the deprived cortex. We then assessed the same parameters in *Cx3cr1*^-/-^ littermates. Consistent with our assessment of structural synapses, remodeling of functional synapses following whisker lesioning was blocked in *Cx3cr1*^-/-^ mice (Fig. 4c,e). There was no longer a significant decrease in sEPSC frequency or amplitude in the deprived cortex compared to the control cortex in *Cx3cr1*^-/-^ mice. Interestingly, there was also a decrease in baseline sEPSC frequency and amplitude in *Cx3cr1*^*-/*-^ mice compared to *Cx3cr1*^+/+^ littermates, suggesting that CX3CR1 signaling may modulate this aspect of functional synapse development within the barrel cortex.

**Figure 4.**
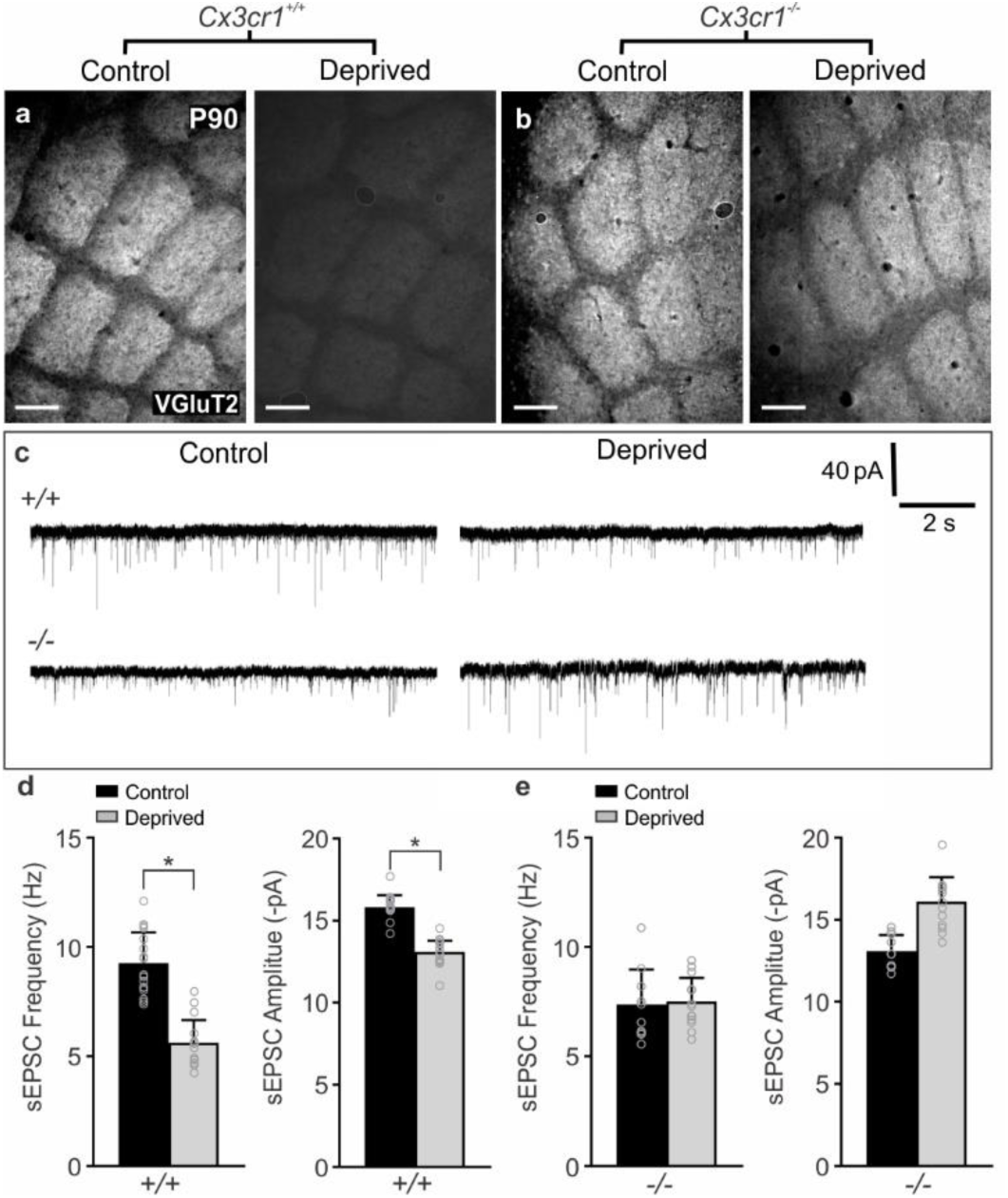
Microglial *Cx3cr1* deficiency blocks structural and functional synaptic remodeling long-term. **a,b**, VGluT2 immunolabeling of TC inputs in tangential sections of the control and deprived barrel cortex in P90 *Cx3cr1*^+/+^ (a) and *Cx3cr1*^-/-^ (b) littermates. TC inputs remain in *Cx3cr1*^-/-^ mice after sustained whisker removal (b, right panel). Scale bar, 150 µm. **c**, Representative sEPSC traces from layer IV stellate neurons for the control and deprived barrel cortices of P42-P56 *Cx3cr1*^+/+^ and *Cx3cr1*^-/-^ mice. **d,e**, Quantification of stellate neuron sEPSC frequency and amplitude in *Cx3cr1*^+/+^ (d; sEPSC Frequency: n = 17 control and 16 deprived cells from 3 *Cx3cr1*^+/+^ littermates, Two-tailed Student’s t-test, *P* = 0.0484, *t* =2.054, *df* =31; sEPSC Amplitude, Two-tailed Student’s t-test, *P* = 0.0105, *t* =2.723, *df* =31) and *Cx3cr1*^-/-^ (e; sEPSC Frequency: n = 10 control and 13 deprived cells from 3 *Cx3cr1*^-/-^ littermates, Two-tailed Student’s t-test, *P* = 0.1286, *t* =1.582, *df* =21; sEPSC Amplitude, Two-tailed Student’s t-test, *P* = 0.9432, *t* =0.07205, *df* =21) mice in the deprived (grey bars) compared to the contralateral control (black bars) barrel cortex. *Cx3cr1*^-/-^ mice show no significant decrease in stellate neuron sEPSC frequency or amplitude. All data presented as mean ± SEM.

This is similar to previously published work showing that CX3CR1 deficiency results in a delay in the functional maturation of synapses^11,12^. These data establish that microglial CX3CR1 signaling is critical for long-term remodeling of structural and functional synapses in the barrel cortex several synapses away from a peripheral sensory lesion.

### CX3CR1 signaling regulates microglia-mediated engulfment of TC synapses

While microglial CX3CR1 signaling has previously been identified to play an important role in regulating synaptic connectivity^11-13^, the mechanism by which CX3CR1 exerts these effects is largely unknown. It has been suggested that CX3CR1, a chemokine receptor, is necessary for microglial recruitment to synapses. Similar to previously published work in the barrel cortex^12^, we found that microglia in neonatal *Cx3cr1*^+/-^ mice were concentrated in barrel septa and infiltrated the barrel centers at P6-7 (Fig. 5a-b and Supplementary Fig. 5). This recruitment to barrel centers was delayed in *Cx3cr1*^-/-^ mice until P8 (Fig. 5a-b). There was no overall difference in the total numbers of microglia in the *Cx3cr1*^-/-^ vs. *Cx3cr1*^+/-^ barrel cortices (Supplementary Fig. 5). We then assessed whether microglial recruitment to barrel centers was affected by whisker lesioning and identified that recruitment and overall density of microglia were largely unaffected in *Cx3cr1*^+/-^ and *Cx3cr1*^-/-^ littermates (Fig. 5a-b, Supplementary Fig. 5). There was still a transient delay in recruitment of microglia to barrel centers in *Cx3cr1*^-/-^ barrel cortex in both control and deprived barrel cortices; however, numbers of microglia within barrel centers were indistinguishable from *Cx3cr1*^+/-^ mice by P8. Given that the bulk of TC synapse elimination occurs after P8 in the current paradigm (Fig. 1a-c), the delay in recruitment of microglia to the barrel centers in <P8 *Cx3cr1*^-/-^ mice likely does not explain the sustained defect in synapse elimination in *Cx3cr1*^-/-^ *mice.*

**Figure 5.**
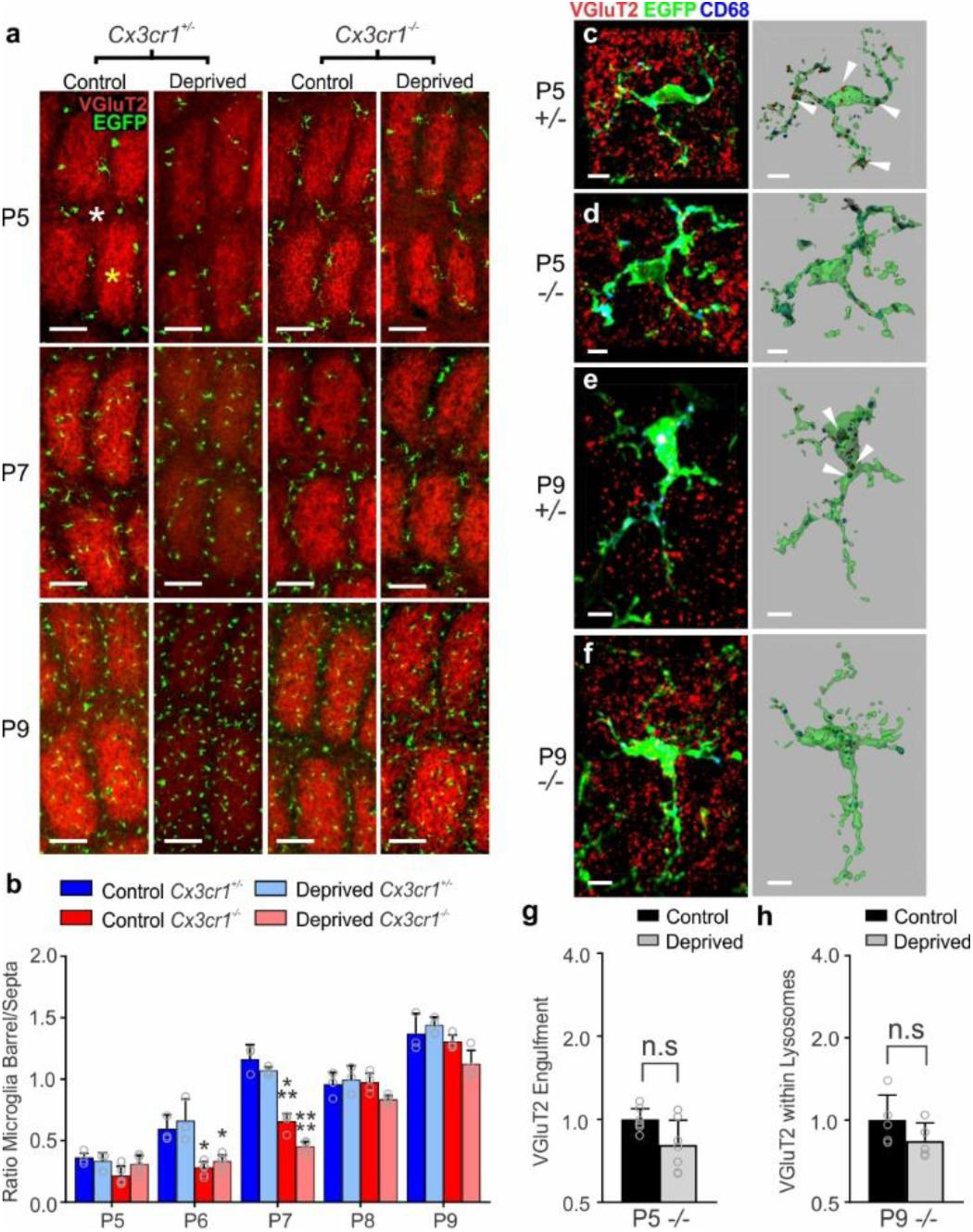
Microglial engulfment of TC inputs following whisker lesioning is CX3CR1-dependent. **a**, Representative images of microglia within the barrel cortex labeled by transgenic expression of EGFP under the control of CX3CR1 (green) and TC inputs labeled with anti-VGlut2 (red). **b**, Quantification of the ratio of microglia localized to the septa (denoted with white asterisk at P5) compared to the barrels (denoted with a yellow asterisk at Pt) +/-whisker deprivation. In both deprived and non-deprived barrel cortices, microglia begin to infiltrate the barrel centers from the septa by P6/7 in *Cx3cr1*^+/-^ mice, which is delayed to P8 in *Cx3cr1*^-/-^ mice. There is no significant difference by P8. (Two-way ANOVA and Tukeys post hoc test, control *Cx3cr1*^+/-^ vs control *Cx3cr1*^-/-^ at P6, n = 3 *Cx3cr1*^+/-^ and 5 *Cx3cr1*^-/-^ littermates, *P* = 0.0394, *q* = 3.887, *df* = 54; deprived *Cx3cr1*^+/-^ vs deprived *Cx3cr1*^-/-^ at P6, n = 3 *Cx3cr1*^+/-^ and 5 *Cx3cr1*^-/-^ littermates, *P* = 0.0273, *q* = 4.092, *df* = 54; *Cx3cr1*^+/-^ vs control *Cx3cr1*^-/-^ at P7, n = 3 *Cx3cr1*^+/-^ and 4 *Cx3cr1*^-/-^ littermates, *P* = 0.0005, *q* = 5.996, *df* = 54; deprived *Cx3cr1*^+/-^ vs deprived *Cx3cr1*^-/-^ at P7, n = 3 *Cx3cr1*^+/-^ and 4 *Cx3cr1*^-/-^ littermates, *P* <0.0001, *q* = 7.32, *df* = 54) **c-f**, Representative microglia from the deprived barrel cortex of *Cx3cr1*^+/-^ and *Cx3cr1*^-/-^ mice 24 hours and 5 d after whisker lesioning. Left panel shows raw fluorescent image with microglia (EGFP, green), VGluT2 (red), and lysosomes (CD-68, blue). Right panel shows 3D-rendered microglia within layer IV of the deprived barrel cortex 24 h (c,d) and 5 d (e,f) post deprivation. Arrows denote examples of engulfed TC inputs in *Cx3cr1*^+/-^ barrel cortex (c,e) which are largely absent in *Cx3cr1*^-/-^ mice (d,f). Scale bar, 5 µm. **g,h**, Quantification of engulfment in *Cx3cr1*^-/-^ mice 24 h (g; Two-tailed Student’s t-test, n = 8 littermate animals, *P =* 0.3668, *t* = 0.938, *df* =12) and 5 d (h; Two-tailed Student’s t-test, n = 5 littermate animals, *P* = 0.5619, *t* =0.6051, *df* =8) post deprivation reveals no significant increase in engulfed TC inputs in the control vs deprived barrel cortex at any time point. Engulfment data normalized to the control, non-deprived hemisphere within each animal. All data presented as mean ± SEM.

To more closely assess microglial-synapse interactions in *Cx3cr1*^-/-^ mice in response to whisker removal, we next analyzed microglia-mediated TC input engulfment. Unlike CR3-KO mice (Fig. 2), loss of CX3CR1 completely blocked TC input engulfment by microglia in the deprived barrel cortex (Fig. 5c-h). This engulfment was blocked at both P5 and P9. Together, these data demonstrate that CX3CR1 signaling modulates microglia-mediated synaptic engulfment and blockade of this engulfment results in sustained defects in structural and functional synapse elimination following a peripheral sensory lesion.

### The CX3CR1 ligand, fractalkine (CX3CL1), is necessary for microglial TC synaptic engulfment and elimination following whisker lesioning

We next sought to understand the signals upstream of CX3CR1 necessary for TC synapse elimination by microglia and first assessed a role for fractalkine (CX3CL1), the canonical ligand of CX3CR1^33^. Unlike CX3CR1, the role of CX3CL1 in microglia-mediated synapse development is largely unknown. Similar to *Cx3cr1*^-/-^ mice, we observed significant defects in the elimination of TC inputs 6 days post-whisker removal in the *Cx3cl1*^-/-^ barrel cortex (Fig. 6a-f). This synapse elimination defect was accompanied by a blockade of TC input engulfment by microglia (Fig. 6g-i) in *Cx3cl1*^-/-^ mice, which phenocopies defects in *Cx3cr1*^-/-^ mice. These data identify CX3CL1 as a novel regulator of TC synapse remodeling and microglia-mediated engulfment of synapses and strongly suggest that microglia eliminate TC inputs through CX3CR1-CX3CL1 signaling.

**Figure 6.**
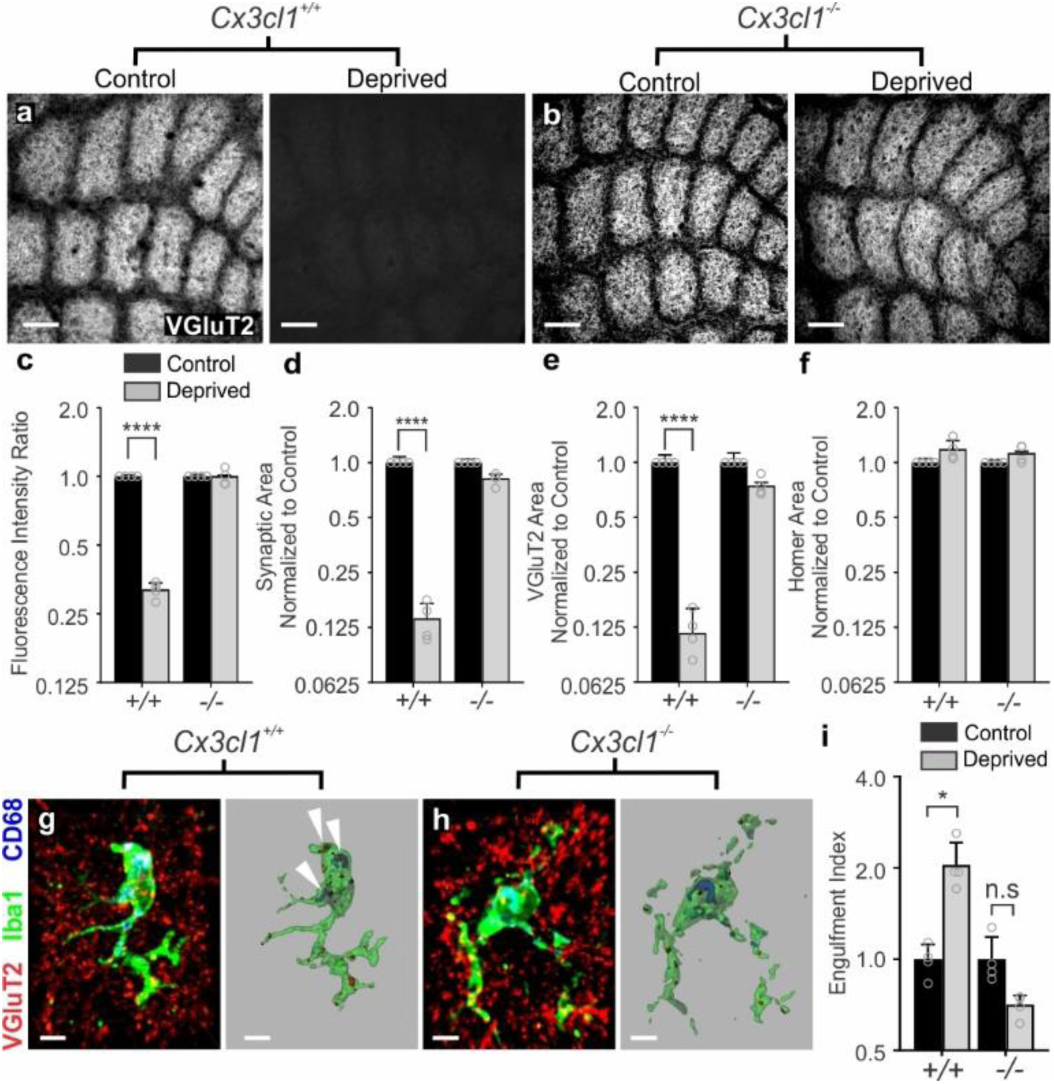
CX3CL1 is necessary for TC input engulfment and elimination after sensory lesioning. **a-b,** VGluT2 immunolabeled TC inputs within tangential sections of control and deprived barrel cortices in *Cx3cl1*^+/+^ (a) and *Cx3cl1*^-/-^ mice (b). Scale bar, 150 µm**. c**, Quantification of fluorescence intensity of VGluT2-positive TC input 6 d after deprivation shows a significant decrease in VGluT2 fluorescence intensity in *Cx3cl1*^+/+^ mice 6 d post-deprivation, which is blocked in *Cx3cl1*^-/-^ littermates. Data normalized to the control, non-deprived hemisphere within each animal. (Two-Way ANOVA with Sidak’s post hoc, n = 4 animals per genotype, *P* <0.0001, *t* = 28.3, *df* = 12). **d-f**, Quantification of high magnification images of synaptic components in the barrel centers 6 d after whisker removal reveals a significant decrease in structural synapses (d, VGluT2 colocalized with Homer; Two-Way ANOVA with Sidak’s post hoc, n = 4 animals per genotype, *Cx3cl1*^+/+^ control vs deprived, *P* <0.0001, *t* =11.66, *df* = 12) and VGluT2-positive presynaptic terminals (e; Two-Way ANOVA with Sidak’s post hoc, n = 4 animals per genotype, *Cx3cl1*^+/+^ control vs deprived, *P* <0.0001, *t* = 7.418, *df* = 12) in *Cx3cl1*^+/+^ mice but no significant change in *Cx3cl1*^-/-^ littermates (Colocalized Area, *P* = 0.0642, t = 2.415, df = 12; VGlut2 Area, *P* = 0.1071, *t* =2.125, *df* = 12). There was no significant change in density of homer immunoreactivity in *Cx3cl1*^+/+^ or *Cx3cl1*^-/-^ mice following whisker deprivation (f; Two-Way ANOVA with Sidak’s post hoc, n = 4 animals per genotype, no significance). Data normalized to the control, non-deprived hemisphere within each animal. **g-h**, Representative microglia from the deprived barrel cortex of *Cx3cl1*^+/+^ (g) and *Cx3cl1*^-/-^ (h) mice. Left panel displays raw fluorescent image with microglia (Anti-Iba1, green) VGluT2 inputs (red) and lysosomes (Anti-CD68, blue) labeled. Right panels shows 3D-surface rendering of these cells. Engulfed VGluT2 (red) immunoreactive TC inputs within microglia are visualized in *Cx3cl1*^+/+^ microglia (g, arrows)) but not *Cx3cl1*^-/-^ microglia (h). Scale bars, 5 µm. **i**, Quantification of VGlut2 engulfment 24 h after whisker removal reveals that *Cx3cl1*^-/-^ microglia fail to engulf TC inputs following sensory deprivation. Data normalized to engulfment in the control hemisphere within each animal. (Two-Way ANOVA with Sidak’s post hoc, n = 4 littermates per genotype, *Cx3cl1*^+/+^ control vs deprived *P* = 0.0111, *t* = 3.369, *df* = 12; *Cx3cl1*^-/-^ control vs deprived *P* = 0.5963, *t* = 0.9422, *df* = 12). All data presented as mean ± SEM.

### Single-cell RNA sequencing reveals that CX3CL1 is highly enriched in neurons in the barrel cortex but its transcription is not modulated by whisker lesioning

To further explore the mechanism by which CX3CL1 signals to CX3CR1 to regulate TC synapse elimination, we performed single-cell RNA sequencing in the deprived and non-deprived barrel cortices of *Cx3cr1*^+/-^ and *Cx3cr1*^-/-^ mice. *Cx3cr1*^+/-^ mice were used as controls as these were the animals used for initial synaptic remodeling and engulfment analyses (Fig. 1). 24 h after unilateral whisker lesioning, when we begin to see TC input engulfment and elimination by microglia (Fig. 1), deprived and control barrel cortices were microdissected from each animal and single-cell RNAseq was performed using inDrops^34^. Following principal component analysis, we identified 27 distinct clusters of CNS cell types within the barrel cortex, which were reproducible across biological replicates with an average read depth across biological replicates of 8,815 reads per cell. (Fig. 7 and Supplementary Figs. 6-7). In agreement with a recent single-cell transcriptomic study in the adult visual cortex identifying changes in gene expression of neuronal and non-neuronal cells in response to visual stimulation, we observed gene expression changes in neurons and glia (Supplementary Figs. 8-9)^35^. To our knowledge, this is the first analysis of sensory lesion and CX3CR1-dependent gene expression in the developing somatosensory cortex at single-cell resolution. From this data set, we identified some potentially interesting differences in glial cell, including microglial, gene expression following whisker lesioning (Supplementary Fig. 9), which were largely blocked in *Cx3cr1*^-/-^ mice (Supplemental Fig. 9a,h). To note, compared to neurons, the numbers of glial cells sequenced was relatively low for individual genotypes (Supplemental Fig. 9g). Therefore, future investigation to validate gene expression in changes in glial cell types is necessary. Going forward, we focused our analyses on whisker lesion-induced changes in gene expression in neuronal populations.

**Figure 7.**
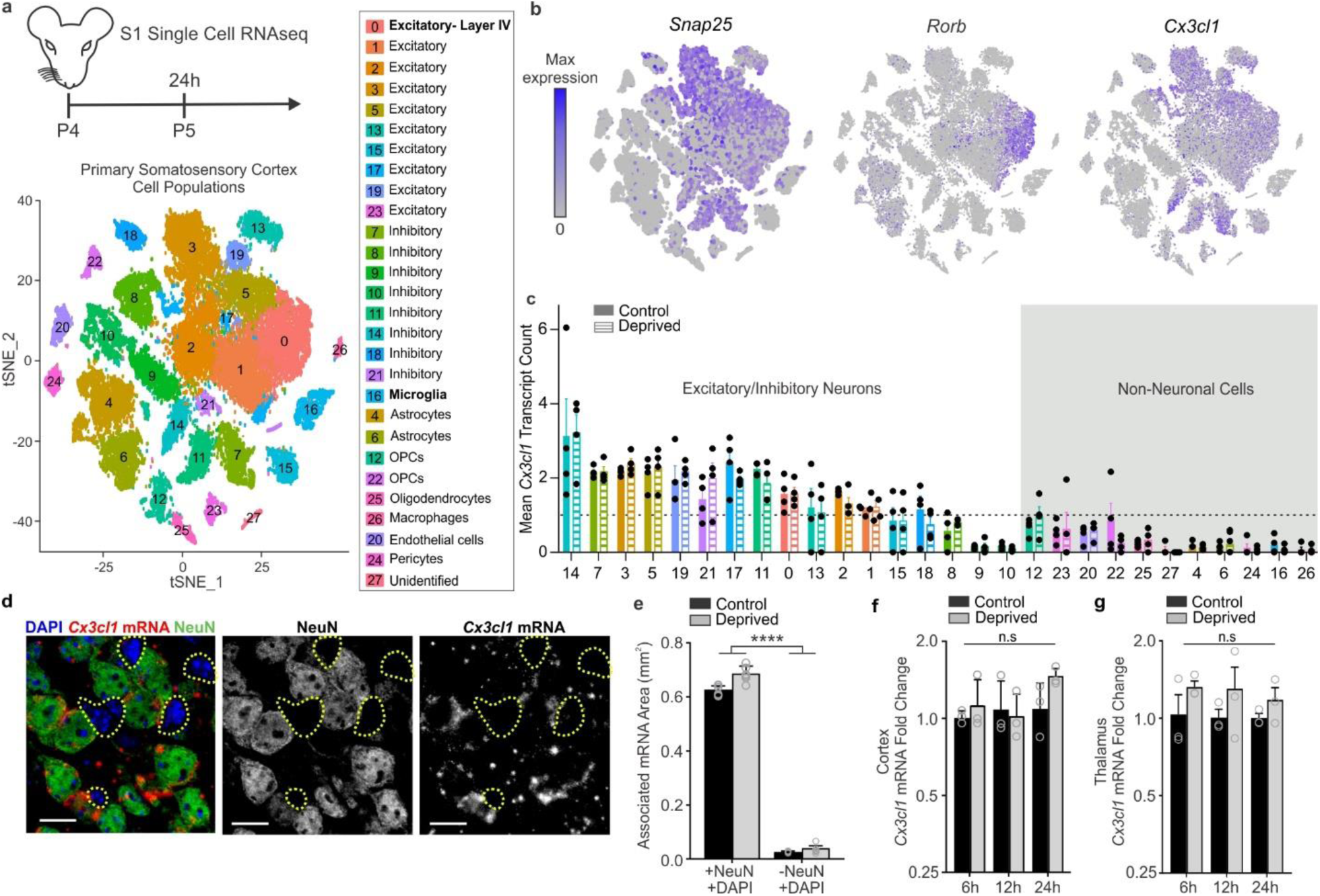
Single-cell RNAseq reveals that *Cx3cl1* is highly enriched in neurons in the barrel cortex but its transcription is not modulated by whisker lesioning. **a**, Timeline for whisker removal and single cell sequencing analysis. P4 mice underwent unilateral whisker lesioning and were sacrificed 24 h later. Barrel cortices were prepared for single-cell RNAseq. A tSNE plot of 27 distinct cell populations in the barrel cortex clustered by principal component analysis. (See Supplementary Fig 7). **b**, tSNE plots for *Snap25* and *Cx3cl1* across all 27 clusters. *Cx3cl1* is enriched in most SNAP-25-poistive neuronal clusters. **c,** Mean *Cx3cl1* RNA transcript counts per condition (control, solid bars; deprived, striped bars). Each data point is the mean for an individual biological replicate. Data at or below the dotted line indicates 1 transcript or no expression. *Cx3cl1* is enriched in neurons but its expression is unchanged following whisker lesioning across all cell types. **d**, *In situ* hybridization for *Cx3cl1* (red) and immunohistochemistry for NeuN to label neurons (green) in *Cx3cl1*^+/+^ deprived barrel cortices validates that *Cx3cl1* is enriched in NeuN-positive neurons compared to non-neuronal cells (NeuN negative, yellow dotted lines). Scale bar, 15 µm. **e**, Quantification of *in situ* for *Cx3cl1* reveals enrichment in neuronal (+NeuN/+DAPI) vs. non-neuronal (-NeuN/+DAPI) cells and no change in expression 24 h-post whisker lesioning. (Two-Way ANOVA with Sidak’s post hoc test, Control +NeuN/+DAPI vs Control -NeuN/+DAPI, *P* <0.0001, *t* = 25.78, *df* = 20; Deprived +NeuN/+DAPI vs Deprived -NeuN/+DAPI, *P* <0.0001, *t* = 27.7, *df* = 27; Control +NeuN/+DAPI vs Deprived -NeuN/+DAPI, *P* <0.0001, *t* = 25.19, *df* = 20; Deprived +NeuN/+DAPI vs Control - NeuN/+DAPI, *P* <0.0001, *t* = 28.29, *df* = 20; n= 6 images from 3 animals). All data presented as mean ± SEM. **f-g,** qPCR for *Cx3cl1* expression in the barrel cortex (f) and VPM nucleus of the thalamus (g) 6, 12, 24, and 72 h after whisker lesioning in *Cx3cr1*^+/+^ mice in the control (black bars) and deprived (grey bars) barrel cortices. (Two-Way ANOVA with Sidak’s post hoc, n = 3 animals per time point, no significant interactions across comparisons).

Similar to previous work^33^, we observed enrichment of *Cx3cl1* mRNA in neurons compared to non-neuronal (Fig. 7b-c). However, despite CX3CL1 being necessary for TC input remodeling following whisker lesioning (Fig. 6), we found no significant difference in *Cx3cl1* mRNA in the deprived vs. control barrel cortex in any cell type (Fig. 7c). These data were confirmed by *in situ* hybridization (Fig. 7d-e). This was further confirmed by RNAseq of whole barrel cortex (Fig 1e) and qPCR from the barrel cortex and VPM nucleus of the thalamus (Fig 7f-g). These data demonstrate that CX3CL1 is neuron-derived, but its transcription in the barrel cortex and thalamus is unchanged following peripheral sensory lesioning.

### ADAM10, a metalloprotease that cleaves CX3CL1, is increased in neurons and microglia within the barrel cortex following whisker lesioning

CX3CL1 can exist in a membrane or a secreted form^33^. The latter is produced upon cleavage of the membrane-bound form by metalloproteases^36^. Therefore, we hypothesized that this post-translational processing of CX3CL1, versus its transcription, was modified upon whisker lesioning. After analyzing gene expression in all 27 cell clusters within our single-cell data set, we identified *Adam10*, a metalloprotease previously shown to be regulated by neuronal activity^37^ and known to cleave CX3CL1^36,38^ (Fig. 8a), as specifically increased in *Rorb*+ layer IV neurons and microglia upon whisker lesioning (Fig. 8b-c and Supplementary Fig. 8). No other cell populations within our single-cell data set showed significant changes in *Adam10* expression. We validated these single-cell data and demonstrated that *Adam10* mRNA increases in *Rorb*+ layer IV neurons in the deprived barrel cortex relative to control by *in situ* hybridization (Fig.8d-e). We then assessed microglial *Adam10* expression by *in situ* in the barrel cortex (Fig. 8g-i). Similar to analysis of *Rorb*+ neurons (Fig. 8e), we averaged the data across all the microglia, but this analysis revealed no significant difference in total microglial *Adam10* expression following whisker lesioning (Fig. 8h). However, we observed a subset of microglia within our images that appeared to have high *Adam10* in the deprived cortex (Fig. 8g). Therefore, we binned the data based on *Adam10* puncta per cell and found that microglia with the highest expression of *Adam10* (>15 puncta/cell) were significantly increased in the deprived vs. control barrel cortex (Fig. 8i). To further validate this upregulation of *Adam10*, we performed qPCR from whole barrel cortex and observed a similar increase in total *Adam10* mRNA 24 h post-whisker lesioning (Fig. 8f). Analysis of bulk RNAseq from whole barrel cortex (Fig. 1d-f) showed a similar effect (Fig 8j-k). These RNAseq data also revealed increases in molecules known to interact with and regulate ADAM10 activity at the membrane (e.g., tetraspanins (*Tspn5 and Tspn14)*; Fig 8k). These data demonstrate that ADAM10, a protease that cleaves CX3CL1 into a secreted form, is induced largely in layer IV excitatory neurons and a subset of microglia within the barrel cortex upon sensory lesioning.

**Figure 8.**
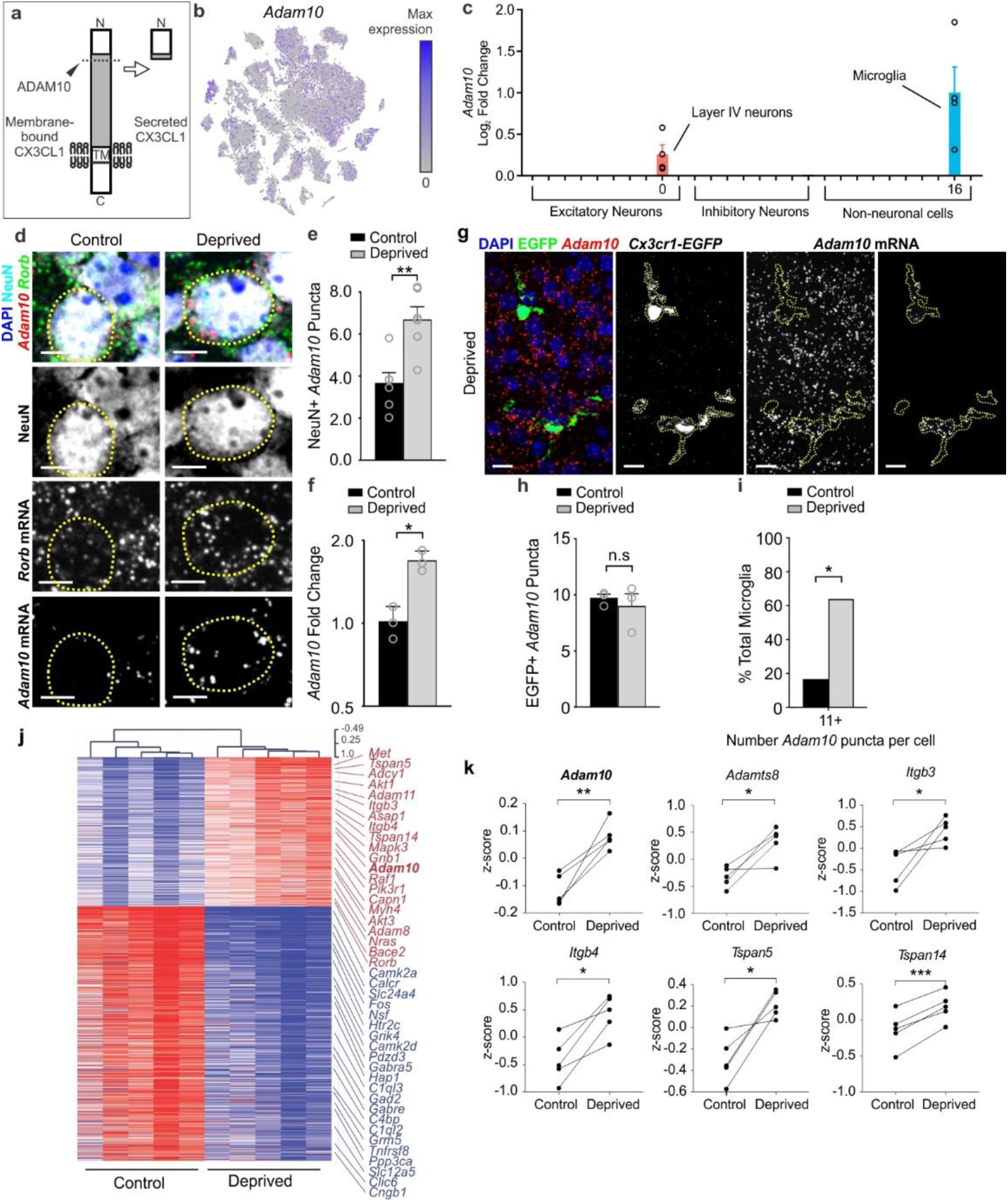
ADAM10, a metalloprotease that cleaves CX3CL1, is increased in neurons within the barrel cortex following whisker lesioning. **a,** ADAM10 cleaves Cx3CL1 at the membrane (dotted line) to produce a secreted form. **b**, tSNE plot for *Adam10* reveals broad expression across many neuronal and non-neuronal cell types. **c**, Fold change of *Adam10* expression by single-cell RNAseq reveals significant (FDR <0.10) upregulation specifically in layer IV Rorb+ neurons and microglia after whisker lesioning. Each data point is an individual biological replicate. **d,** *In situ* hybridization for *Adam10* in the control (top panels) and deprived (bottom panels) barrel cortices. *Adam10* is increased in the majority of *Rorb*+ layer IV excitatory neurons (NeuN+, *Rorb*+) assessed 24 h-post whisker lesioning compared to neurons in the control barrel cortex. Scale bar, 5 µm. **e,** Quantification of *in situ* for *Adam10* puncta co-localized with layer IV neurons. (Two-tailed Student’s t-test, n = 6 *Cx3cr1*^+/+^ animals, *P =* 0.003, *t* = 0.3889, *df* =10). **f**, qPCR for *Adam10* 24 h-post whisker lesioning in the control (black bars) and deprived (grey bars) whole barrel cortices reveals a significant increase in Adam 10 24 h-post whisker lesioning. (Two-tailed Student’s t-test, n = 3 *Cx3cr1*^+/+^ animals, *P =* 0.0241, *t* = 3.538, *df* =4). **g**, *In situ* hybridization for *Adam10* within *Cx3cr1*^*EGFP/+*^ microglia (yellow dotted lines) in the deprived cortex 24 h-post whisker lesioning reveals increased *Adam10* expression in a subset of microglia after lesioning. **h**, Quantification of the average *Adam10 in situ* puncta per microglia averaged across all microglial cells assessed in the barrel cortex shows no significant difference in expression between the control and deprived conditions. (Two-tailed Student’s t-test, n = 3 *Cx3cr1*^+/-^ animals, *P =* 0.5547, *t* = 0.6439, *df* =4). **i**, Further quantification of *Adam10* mRNA puncta within microglia reveals a significant increase in a subset of microglia expressing high levels (≥11 puncta) of *Adam10* in the deprived vs. control barrel cortex (One-tailed Chi-square test, *P* = 0.0306, χ^2^ = 3.503, *df* = 1, *z* = 1.872). **j**, Heatmap with hierarchical clustering distances shows the variation in the expression levels (z-scored log2(RPKM)) of 539 up- and 918 downregulated genes upon whisker deprivation at P4 identified by bulk RNAseq from the primary barrel cortex of whisker lesioned mice (DESeq2 software, n = 5 mice, P5, related to Figure1e). *Adam10* is bolded. **k,** Line graphs show for individual mice z-scored log2(RPKM) changes of *Adam10* and other selected genes encoding known regulators of ADAM10 expression or activity. (Two-tailed Student’s t-test: *Adam10, P* = 0.0060, *t* = 5.32; *Adamts8, P* = 0.0205, *t* =3.72; *Itgb3, P* = 0.0454, *t* = 2.87; *Itgb4, P* = 0.0106, *t* = 4.53; *Tspan5, P* = 0.0151, *t* = 4.08; *Tspan14, P* = 0.0006, *t* = 9.80; n = 5 animals).

### Pharmacological inactivation of ADAM10 phenocopies TC synapse elimination defects in *Cx3cr1*^-/-^ and *Cx3cl1*^-/-^ mice

To then determine if induction of *Adam10* in response to whisker lesioning translated to circuit level changes in TC synapse remodeling as observed in *Cx3cr1*^-/-^ and *Cx3cl1*^-/-^ mice, we pharmacologically inhibited ADAM10. Whiskers were lesioned at P4 and a pharmacological inhibitor of ADAM10 (GI254023X, 25 mg/kg), which has been previously been demonstrated *in vivo* to cross the blood brain barrier and specifically inhibit ADAM10 vs. other ADAMs^39-41^, was administered intraperitoneally (IP) daily starting at the time of whisker lesioning (Fig. 9a). TC synapse engulfment and elimination were subsequently assessed in the control and deprived barrel cortices. Identical to *Cx3cr1*^-/-^ and *Cx3cl1*^-/-^ mice, daily administration of the ADAM10 inhibitor (25mg/kg) by IP injection (P4-P10) resulted in profound disruption of TC input elimination following whisker cauterization (Fig. 9b-c). This was concomitant with a blockade of microglia-mediated engulfment of TC inputs (Fig. 9d-i). These data suggest a mechanism by which post-translational modification of CX3CL1 by ADAM10 is increased in neurons following sensory lesioning, which is known to decrease activity in the barrel cortex. This secreted CX3CL1 then signals to microglia via CX3CR1 to eliminate synapses.

**Figure 9.**
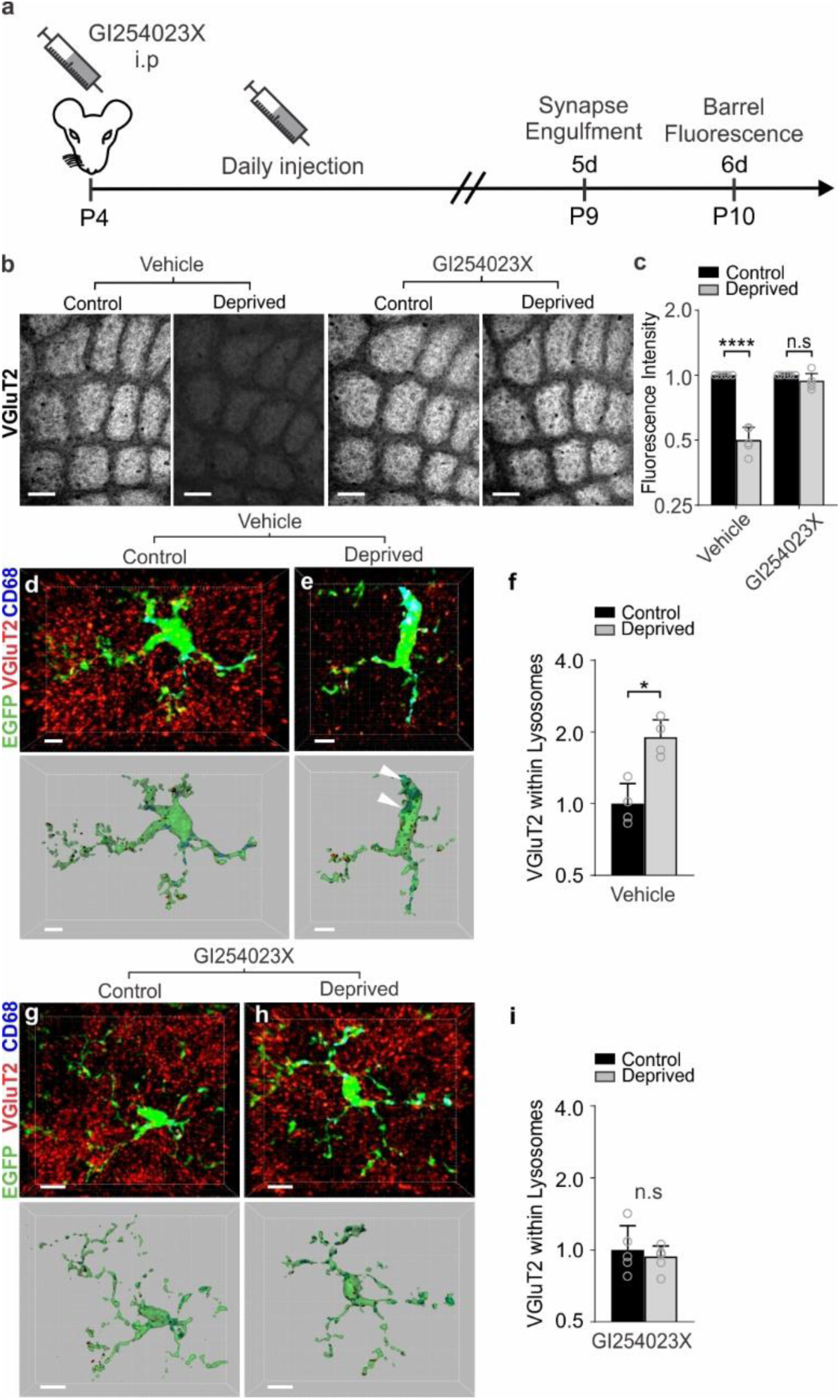
Pharmacological inactivation of ADAM10 phenocopies TC synapse elimination defects in *Cx3cr1*^-/-^ and *Cx3cl1*^-/-^ mice. **a**, Timeline for pharmacological inhibition of ADAM10 via daily 25 mg/kg GI254023X injections intraperitoneally. **b**, Inhibition of ADAM10 (right panels), but not vehicle treatment (left panels), blocks TC input loss as visualized by immunostaining for VGluT2. Scale bars, 150 µm. **c,** Quantification of VGluT2 immunostaining intensity 5 days-post whisker lesioning and GI254023X injections (Two-Way ANOVA with Sidak’s post hoc, n = 5 *Cx3cr1*^+/+^*;Cx3cl1*^+/+^ animals per condition, Vehicle control vs deprived, *P* <0.0001, *t* = 6.782, *df* = 16; GI254023X control vs deprived, P = 0.9715, *t* = 0.7789, *df* = 16). **d,e,** Representative microglia from the control (d) and deprived (e) cortices of vehicle treated *Cx3cr1*^*EGFP/+*^ mice. Top panel displays raw fluorescent image with microglia (EGFP, green) VGluT2 inputs (red) and lysosomes (Anti-CD68, blue) labeled. Bottom panels shows 3D-surface rendering of these cells. Engulfed VGluT2 (red) immunoreactive TC inputs within microglia are visualized in *Cx3cr1*^*EGFP/+*^ microglia (e, arrows)) in the deprived cortex but not the control cortex (d). Scale bars, 5 µm. **f**, Quantification of engulfed VGlut2 5 days after whisker lesioning reveals increased engulfment in the deprived cortex of vehicle treated mice. Data normalized to engulfment in the control hemisphere within each animal. (One-tailed Student’s T-test, n = 4 *Cx3cr1*^*EGFP/+*^ mice, control vs deprived *P* = 0.0455, *t* = 2.012, *df* = 6.) **g,h** Representative microglia from the control (d) and deprived (e) cortex of GI254023X treated *Cx3cr1*^*EGFP/+*^ mice 5 days after whisker lesioning. Scale bars, 5 µm. **i**, Quantification of engulfment 5 days-post whisker lesioning in GI254023X treated mice reveals a blockade of engulfment following ADAM10 inhibition. Data normalized to engulfment in the control hemisphere within each animal. (Two-tailed Student’s T-test, n = 4 *Cx3cr1*^*EGFP/+*^ mice, control vs deprived *P* = 0.8291, *t* = 0.2231, *df* = 6). All data presented as mean ± SEM.

## DISCUSSION

We have identified a novel mechanism by which neurons signal to microglia to remodel synapses in response to a changing sensory environment. Following whisker lesioning, microglia engulf TC inputs in the barrel cortex, which is blocked in CX3CR1 and CX3CL1-deficient mice. Importantly, loss of CX3CR1 signaling, but not CR3, results in sustained defects in the elimination of structural and functional synapses following whisker lesioning. Using single cell RNAseq analysis, we then identify that *Cx3cl1* is expressed primarily by neurons. Moreover, *Adam10*, a gene encoding a metalloprotease known to cleave CX3CL1 into a secreted form, is specifically induced in layer IV excitatory neurons and a subset of microglia upon sensory lesioning. As the first single-cell transcriptomic study in the developing barrel cortex, this is a rich dataset and likely a valuable resource for assessing gene expression in glia and neurons. As validation of one gene hit, we demonstrate using 3 additional methods that *Adam10* mRNA is increased upon whisker lesioning, the mRNA of known regulators of ADAM10 are also modulated following whisker lesioning, and pharmacological inhibition of ADAM10 phenocopies TC synapse elimination defects observed in CX3CR1 and CX3CL1-deficient mice. Together, our data suggest a mechanism by which post-translational processing of CX3CL1 in neurons by ADAM10 is regulated by changes in network activity following a peripheral sensory lesion and subsequently cleaved CX3CL1 signals to microglia to engulf and eliminate synapses via a CX3CR1-dependent mechanism. These data suggest an active process of microglia-mediated TC input elimination as blocking engulfment results in defects in sustained increases in structural and functional synapses. Importantly, we have identified a novel mechanism of neuron-microglia communication that, upon a peripheral sensory lesion several synapses away, leads to the remodeling and elimination of thalamocortical synapses.

Published studies have identified that microglia contact and/or engulf synaptic elements and these interactions are activity-dependent^1,2,29,42-49^. However, the mechanisms underlying these activity-dependent microglia-synapse interactions and the neuron-derived signals driving this process remain largely unknown. Another transcriptional profiling study has demonstrated that, under basal conditions, microglia express a number of transcripts for sensing endogenous ligands and microbes, which was termed the ‘sensome^50^.’ Our data provide a role for one key microglial ‘sensome’ gene *Cx3cr1* in microglia-mediated synapse elimination following manipulation of circuit activity via whisker lesioning or trimming. This is consistent with previous work demonstrating roles for CX3CR1 in modulating microglial function at synapses during development and after seizures^11-13 49^. Our work provides important mechanistic insight into how CX3CR1 is regulating microglial function at synapses and identifies a new role for neuron-derived CX3CL1 and ADAM10 in this process. Going forward, it will be important to elucidate precisely how ADAM10 is modulated in neurons and microglia following sensory lesioning. Also, developing new tools to monitor CX3CL1 cleavage in specific cells will be another important future direction.

This study also raises a new question regarding how downstream CX3CR1 signaling is regulating microglial engulfment and synapse elimination. Our single-cell data set provide some evidence that there is a downregulation of genes related to phagocytosis in *Cx3cr1*^-/-^ microglia, even without whisker lesioning (Supplementary Fig. 10). This raises the possibility that CX3CR1 signaling could modulate basal phagocytic state of microglia through changes in gene expression. In addition, by our single-cell analyses, microglia, as well as other glia, appear to modify their gene expression in response to sensory lesioning. While these gene expression changes require further validation, this is an exciting first step in understanding how microglia respond to a changing sensory environment. We have evidence that these lesion-induced microglial gene expression changes are dramatically modified by loss of CX3CR1. Therefore, another exciting future direction is to further elucidate these CX3CR1-dependent gene expression changes.

Another important theme that emerges from our data is that different neural-immune signaling mechanisms are utilized by microglia to engulf and remodel synaptic connections and these mechanisms appear to be engaged in a context-dependent manner. One recent study in the developing rodent spinal cord has shown a role for astrocytic IL-33 in mediating microglial synapse engulfment through IL1RL1^51^. Work in the developing mouse retinogeniculate identified complement-dependent phagocytic signaling as a key regulator of microglial engulfment and elimination of synapses undergoing developmental pruning^1,6,7^ and microglial P2RY12 was identified as a key regulator of ocular dominance plasticity in the visual cortex^29^. These findings are in line with seminal work discovering that MHC class I molecules, another immune pathway, are critical regulators of synapse development and plasticity in the visual system^52-54^. Interestingly, retinogeniculate pruning and ocular dominance plasticity occur independent of CX3CR1^9,10^. In contrast, we show that sensory deprivation-induced elimination of synapses in the barrel cortex is regulated by CX3CR1-CX3CL1 signaling, but not CR3. These data demonstrate that distinct immune mechanisms are utilized to regulate synaptic remodeling depending on the circuit and/or paradigm used for manipulating activity. For example, the whisker lesioning paradigm used here induces injury to the peripheral sensory endings in the snout several synapses away from the barrel cortex. Therefore, this may be an injury vs. activity-dependent response. However, there are several lines of evidence arguing against this being an injury response. First, there is no change in neuronal cell death or degeneration following whisker lesioning. Second, there is a similar, albeit delayed, CX3CR1-depenedent synapse elimination following whisker trimming. Last, our RNAseq data demonstrate there is a downregulation in genes related to neural activity, but, besides genes related to phagocytosis, no genes associated with injury or inflammation. These data are most consistent with cortical activity modulating CX3CR1-CX3CL1 signaling via ADAM10 and microglia-mediated synapse elimination.

A broad range of neurological disorders from autism and schizophrenia to traumatic injuries causing loss of eye sight, cutaneous sensation, olfaction or hearing all result in changes in sensory inputs and TC connectivity^55-60^. Further, defects in CX3CR1-CX3CL1 signaling and ADAM10 have been identified to either enhance or suppress neurodegeneration in a variety of neurological disease models depending on the disease, insult, brain region, etc.^38,61-66^. Here, we identify that neuronal ADAM10 is modulated in the cortex by sensory lesioning and disruption of ADAM10, CX3CL1 (an ADAM10 substrate), or CX3CR1 results in profound defects in microglial synaptic engulfment and synapse elimination following a peripheral sensory lesion known to dampen activity in the cortex^38,64-66^. This mechanism is particularly intriguing in light of recent data in mouse models of Alzheimer’s disease where changes in neural activity modulates microglial morphology, Aβ, and amyloid plaques^67,68^. One exciting possibility is that activity dependent CX3CR1-CX3CL1-ADAM10 signaling could be involved in modulating plaque pathology via microglia. Together, our findings have strong implications for advancing our understanding of how microglia and CX3CR1-CX3CL1-ADAM10 signaling affect brain circuits and provide novel insight into how TC circuits remodel in the cortex. These insights are important for our basic understanding of how neurons communicate with microglia to modulate neural circuits and have significant translational potential for a variety of neurological disorders with underlying changes in synaptic connectivity, microglia, and fractalkine signaling.

## Acknowledgments

We thank M. Freeman (OHSU), V. Budnik (UMMS), E. Baehrecke (UMMS), M. Francis (UMMS), P. Greer (UMMS), and R. Bruno (Columbia University) for critical reading of the manuscript. The *Cx3cl1*^-/-^ mice and SERT-Cre mice were generously provided by S. Lira (MSSM) and M. Ansorage/S. Nelson (Columbia University/Brandeis University), respectively. We thank M. Cahill (UMMS) and A. Lotun (UMMS) for assistance assessing microglia within the barrel cortex and S. Becker (UMMS) and J. Jung (UMMS) for assistance with tissue preparation and whisker trimming experiments. H. Learnard (UMMS), A. Song (UMMS), Z. Zhang (Boston Children’s Hospital), and C. Woolf (Boston Children’s Hospital) for assistance with experiments to assess ATF3. This work was funded by NIMH-R00MH102351 (DPS), NIMH-R01MH113743 (DPS), NIMH-R21MH115353 (DPS), Charles H. Hood Foundation (DPS), Brain & Behavior Research Foundation (DPS), Worcester Foundation (DPS), and the Dr. Miriam and Sheldon G. Adelson Medical Research Foundation (DPS).

## Author Contributions

G.G. and D.P.S. designed the study, performed most experiments, analyzed most data, and wrote the manuscript, K.J. assisted in the design of initial experiments and performed experiments to identify initial synapse remodeling and engulfment phenotypes, L.C., A.N., and M.E.G. performed single-cell sequencing experiments. E.M. performed in situ hybridization experiments, P.A., A.B., and A.S. performed bulk RNAseq experiments of whole barrel cortex, L.L. and A.R.T. performed electrophysiology experiments. K.K., S.M.B. and B.T.L. performed experiments related to *Cx3cl1*^-/-^ mice, R.M.R. provided critical input into study design and feedback on writing of the manuscript.

## Declaration of Interests

Authors declare no conflict of interest.

## Materials and Methods

### Animals

SERT-Cre mice were a generous gift from Dr. Mark Ansorage, Columbia University and provided by Dr. Sacha Nelson, Brandeis University. *Cx3cl1*^-/-^ mice were provided by Dr. Sergio Lira (Ichan School of Medicine, Mount Sinai). Rosa26-TdTomato mice (Ai14; stock #007914), Cx3cr1^-/-^ mice (Cx3cr1^EGFP/EGFP^; stock #005582), CR3-KO mice (stock #003991), and C57Bl6/J (stock #000664) mice were obtained from Jackson Laboratories (Bar Harbor, ME). Heterozygous breeder pairs were set up for all experiments and wild-type and heterozygote littermates were used as controls with equal representation of males and females for each genotype. All experiments were performed in accordance with animal care and use committees and under NIH guidelines for proper animal welfare.

### Whisker Removal

For sensory lesioning, whiskers were unilaterally removed at P4 with a high temperature handheld cautery kit (Bovie Medical Corporation; Clearwater, FL) applied to the right whisker pad of anesthetized pups. After whisker removal, pups were placed on a heating pad before animals were returned to their home cage. For long term whisker deprivation, whiskers were removed at P4 and assessed for whisker re-growth weekly. For whisker trimming experiments, whiskers were unilaterally trimmed from the right whisker pad starting at P4. Whiskers were trimmed twice daily to minimize whisker growth up to P21.

### Immunohistochemistry

Animals were perfused with 4% paraformaldehyde in 0.1M phosphate buffer (PB) prior to brain removal. In order to visualize the entire barrel field in one plane, the midbrain was dissected from each brain hemisphere and the cortex was then flattened between two slides in 4% paraformaldehyde overnight. Sections were placed in 30% sucrose in 0.1M PB for 24 hours before 50µm (P5-P11) and 40µm (>P40) tangential sections were prepared. Sections were blocked in 10% goat serum, 0.01% TritonX-100 in 0.1M PB for 1 hour before primary immunostaining antibodies were applied overnight. For analysis of synaptic engulfment within microglia, anti-CD68 (AbD Serotec; Raleigh, NC) and anti-VGluT2 (MilliporeSigma; Darmstadt, Germany) were used. Microglia were labeled either using transgenic expression of EGFP (CX3CR1^EGFP/+^) or immunostaining for anti-Iba-1 (Wako Chemicals; Richmond, VA). For analysis of synapse density, anti-VGluT2 (MilliporeSigma; Darmstadt, Germany) and anti-Homer1 (Synaptic Systems; Goettingen, Germany) were used. For markers of neurodegeneration, cell death, and cell stress, anti-APP (ThermoFisher Scientific; Waltham, MA), anti-Cleaved Caspase 3 (Cell Signaling Technology; Danvers, MA), and anti-ATF3 (Sigma-Aldrich; Darmstadt, Germany) were used. Anti-NeuN (MilliporeSigma; Darmstadt, Germany) was used as a marker for neuronal cell bodies. Peripheral monocytes/macrophages were labelled with anti-CD45 (Bio-Rad; Hercules, CA).

### Fluorescence Intensity Analysis

For fluorescence intensity, single plane 10x epifluorescence images were collected at the same exposure time with a Zeiss Observer microscope equipped with Zen Blue acquisition software (Zeiss; Oberkochen, Germany). Fluorescence intensity within each barrel was quantified similar to what has been described previously^22^. Briefly, each image was analyzed in ImageJ (NIH) where first all image pixel intensity thresholds were set to the full range of 16-bit images before quantification to ensure a consistent pixel range across all images. To sample fluorescence intensity, a circular region of interest (ROI) 75µm in circumference (4470.05 µm^2^ area) was placed within the center of 15-20 barrels per 10x field of view. A background ROI outside of the barrel field was also taken for each image. The raw integrated density of pixels within each ROI was measured and each barrel intensity value was background-corrected by subtraction of the background ROI pixel intensity. Average intensity over all barrel ROIs was quantified for each image and then normalized to the control hemisphere within each animal. All data analyses were performed blind to genotype.

### Engulfment Analysis

Engulfment analysis was performed according to previously described methods^1,31^. Briefly, immunostained sections were imaged on a Zeiss Observer Spinning Disk Confocal microscope equipped with diode lasers (405nm, 488nm, 594nm, 647nm) and Zen acquisition software (Zeiss; Oberkochen, Germany). For each hemisphere, three to five 63x fields of view within the barrel field were acquired with 50-70 z-stack steps at 0.27 µm spacing. Images were first processed in ImageJ (NIH) and then individual images of 15-20 single cells per hemisphere per animal were processed in Imaris (Bitplane; Zurich, Switzerland) as previously described. All image files were blinded for unbiased quantification. All data was then normalized to the control, spared hemisphere within each animal. Note, *Cx3cr1*^+/-^ littermates were used for comparison to *Cx3cr1*^-/-^ mice as microglia within both sets of mice are labeled with EGFP and show similar changes in TC synapses (Fig 2).

### Synapse Density Analysis

Synapse density analysis was performed blind to condition and genotype as described previously ^*1,3*^. Briefly, immunostained sections were imaged on a Zeiss LSM700 scanning confocal microscope equipped with 405nm, 488nm, 555nm, and 639nm lasers and Zen acquisition software (Zeiss; Oberkochen, Germany). Synapse density analysis was performed on single plane confocal images in ImageJ (NIH). Three 63x fields of view per hemisphere per animal were analyzed. Sample images for each genotype and condition were manually thresholded by eye, and a consistent threshold range was determined (IsoData segmentation method, 85-255). Each channel was thresholded and the Analyze Particles function with set parameters for each marker (VGluT2 = 0.2-infinity; Homer1 = 0.1-infinity) was used to measure the total pre- and postsynaptic puncta area. To quantify total synaptic area, Image Calculator was used to visualize co-localized pre- and postsynaptic puncta and then the Analyze Particles function was used to calculate the total area of co-localized puncta. Data for each hemisphere was averaged across all three fields of view and then normalized to the control hemisphere within each animal.

### Bulk RNA sequencing

Mice were killed by CO2 asphyxiation at indicated ages, and brain regions of interest were dissected. Brain tissue from one mouse was immediately homogenized with a motor-driven Teflon glass homogenizer in ice-cold polysome extraction buffer (10 mM HEPES (pH 7.3), 150 mM KCl, 5 mM MgCl_2_, 0.5 mM dithiothreitol (Sigma) 100 μg/mL cycloheximide (Sigma), EDTA-free protease inhibitor cocktail (Roche), 10 μL/mL RNasin (Promega) and Superasin (Applied Biosystems). Homogenates were centrifuged for 10 min at 2,000g, 4°C, to pellet large cell debris. NP-40 (EMD Biosciences, CA) and 1,2-diheptanoyl-sn-glycero-3-phosphocholine (Avanti Polar Lipids, AL) were added to the supernatant at final concentrations of 1% and 30 mM, respectively. After incubation on ice for 5 min, the lysate was centrifuged for 10 min at 13,000g to pellet insoluble material. RNA was purified from the lysate using RNeasy Mini Kit (Qiagen) following the manufacturer’s instructions. RNA integrity was assayed using an RNA Pico chip on a Bioanalyzer 2100 (Agilent, Santa Clara, CA), and only samples with RIN > 9 were considered for subsequent analysis. Double-stranded cDNA was generated from 1–5 ng of RNA using Nugen Ovation V2 kit (NuGEN, San Carlos, CA) following the manufacturer’s instructions. Fragments of 200 bp were obtained by sonicating 500 ng of cDNA per sample using the Covaris-S2 system (duty cycle: 10%, intensity: 5.0, bursts per second: 200, duration: 120 s, mode: frequency sweeping, power: 23 W, temperature: 5.5–6°C; Covaris Inc., Woburn, MA). Subsequently, these fragments were used to produce libraries for sequencing by TruSeq DNA Sample kit (Illumina, San Diego, CA, USA) following the manufacturer’s instructions. The quality of the libraries was assessed by 2200 TapeStation (Agilent). Multiplexed libraries were directly loaded on NextSeq 500 (Illumina) with high-output single-read sequencing for 75 cycles. Raw sequencing data was processed using Illumina bcl2fastq2 Conversion Software v2.17. Raw sequencing reads were mapped to the mouse genome (mm9) using the TopHat2 package (v2.1.0). Reads were counted using HTSeq-count (v0.6.0) against the Ensembl v67 annotation. The read alignment, read count, and quality assessment using metrics, such as total mapping rate and mitochondrial and ribosomal mapping rates, were done in parallel using an in-house workflow pipeline called SPEctRA. The raw counts were processed through a variance stabilizing transformation (VST) procedure using the DESeq2 package to obtain transformed values that are more suitable than the raw read counts for certain data mining tasks. Principal component analysis (PCA) was performed on the top 500 most-variable genes across all samples based on the VST data to visually assess whether there were any outliers. Additionally, hierarchical clustering was used to assess the outliers once again to protect against false positives/negatives from PCA, and the outliers were further justified by the aforementioned quality control metrics as well as experimental metadata. After outlier removal, all pairwise comparisons were performed on the count data of entire gene transcripts using the DESeq2 package (v1.6.3). A cutoff of adjusted P value < 0.05, and mean expression > 3.5. MA plot was made using R (v3.1.1; https://www.R-project.org). For the heatmap, expression of each gene in log2(RPKM) was normalized to the mean across all samples (*z*-scored). Heatmap with hierarchical clustering was made on Multiple Experiment Viewer 4.8 (v.10.2; http://www.tm4.org/) with Pearson correlation by average link clustering. Bar graphs representing RPKM of genes and line plots representing z-scored log2(RPKM) values were made on GraphPad Prism v5.01 (https://www.graphpad.com).

### Single-cell transcriptomics experimental design

Non-deprived, control and deprived barrel cortices from a total of 8 mice served as the input for single-cell sequencing studies, equaling 16 samples total. The capture and barcoding of single cells was performed across four separate experiments on two different days. Per experiment, one littermate each of the genotypes Cx3cr1^+/-^ and Cx3cr1^-/-^ underwent unilateral whisker cauterization at P4 and their deprived and non-deprived cortices were then processed in parallel at P5, yielding 4 samples per experiment. 3,000 cells were collected from each hemisphere of each mouse, equaling a total of ∼48,000 cells collected overall. All libraries were prepared in parallel then pooled and sequenced together across two sequencing runs, as described below.

### Generation of single-cell suspensions

Single-cell sequencing of mouse cortex using inDrops was performed as previously described^35,^. Mice were transcardially perfused with ice-cold choline solution containing 2.1 g NaHCO_3_ per liter, 2.16 g glucose per liter, 0.172 g NaH_2_PO_4_·H_2_O per liter, 7.5 mM MgCl_2_·6H_2_O, 2.5 mM KCl, 10 mM HEPES, 15.36 g choline chloride per liter, 2.3 g ascorbic acid per liter, and 0.34 g pyruvic acid per liter (all chemicals, Sigma). A caveat of sequencing techniques that involve mechanical and enzymatic dissociation is that the dissociation itself induces neural activity-dependent and injury-induced gene transcription^35^. Therefore, because we were interested in analyzing sensory experience-dependent changes in gene expression across all cell types, a number of drugs that block neuronal activity and transcription were included in the perfusion solution. These included 1 μM TTX (Sigma), 100 μM AP-V (Thermo Fisher Scientific), 5 μg actinomycin D (Sigma) per milliliter, and 10 μM triptolide (Sigma). Following the 5 minute perfusion, deprived and non-deprived, control somatosensory cortices were microdissected and each sample was transferred to a tube with 1.65 mL of pre-incubation solution containing HBSS (Life Technologies), 10 mM HEPES, 172 mg kynurenic acid (Sigma) per liter, 0.86 g MgCl_2_·6H_2_O per liter, and 6.3 g D-glucose (Sigma) per liter, pH 7.35, which was saturated with 95% O_2_ and 5% CO_2_ previously but was not bubbled after the sample was added. The drugs contained in the perfusion solution were also present in the pre-incubation solution at the same concentrations. After 30 minutes on ice, 1.65 mL of papain (Worthington) was added to a final concentration of 20 U per milliliter. Samples were moved to a rocker at 37° C and gently rocked for 60 minutes.

Following the papain incubation at 37° C, a series of triturations in increasingly small volumes were performed to fully dissociate the tissue. In between each round of trituration, the tissue was filtered through the corner of a 40 μM nylon cell strainer (Corning). The cells were then centrifuged at 300*g for 5 minutes and the pellet resuspended in 1 mL of trypsin inhibitor (Worthington) plus DNAse (Sigma) in preincubation solution without drugs (dissociation media, DM). The cells were washed by resuspension in DM adjusted to 0.04% BSA (Sigma) three times, then resuspended in DM containing 0.04% BSA and 15% Optiprep (Sigma) to a concentration of 100,000 cells per mL and transferred to the Single-Cell Core at Harvard Medical School for inDrops collection (see Acknowledgements).

### Single-cell RNA sequencing via inDrops

For each sample, approximately 3,000 cells were encapsulated into microfluidic droplets containing polyacrylamide gels with embedded barcoded reverse transcription primers. Reverse transcription was carried out in intact droplets to generate barcoded cDNA from a single cell. Following droplet lysis, inDrops libraries were prepared as previously described^34, 69^. All 16 libraries were indexed, pooled, and sequenced (Read 1: 54 cycles, Read 2: 21 cycles, Index 1: 8 cycles, Index 2: 8 cycles) across 2 runs on a NextSeq 500 (Illumina) with an average read depth across biological replicates of 8,815 reads per cell.

### inDrops data processing

Sequenced reads were processed according to a previously published pipeline^70^. Briefly, this pipeline was used to build a custom transcriptome from Ensembl GRCm38 genome and GRCm38.84 annotation using Bowtie 1.1.1, after filtering the annotation gtf file (gencode.v17.annotation.gtf filtered for feature_type=”gene”, gene_type=“protein_coding” and gene_status=“KNOWN”). Read quality control and mapping against this transcriptome were performed. Unique molecular identifiers (UMIs) were used to link sequence reads back to individual captured molecules. All steps of the pipeline were run using default parameters unless explicitly stated.

### Quality control and clustering of cells

All cells were combined into a single dataset. Nuclei with >10% mitochondrial content were excluded from the dataset. Cells with fewer than 400 UMI counts were excluded. Cells were then clustered using the Seurat R package^71^. The data were log normalized and scaled to 10,000 transcripts per cell. Variable genes were identified using the following parameters: x.low.cutoff = 0.0125, x.high.cutoff = 3, y.cutoff = 0.5. We limited the analysis to the top 30 principal components. Clustering resolution was set to 0.6. The expression of known marker genes was used to assign each cluster to one of the main cell types. *Tubb3*/*Snap25* was used to identify neurons, *Gad1/Gad2* was used to identify inhibitory neurons, *Rorb* was used to identify excitatory cortical layer IV neurons, *Aldoc*/*Aqp4* was used to identify astrocytes, *Mbp*/*Plp1* was used to identify mature oligodendrocytes, *Pdgfra*/*Matn4* was used to identify immature oligodendrocytes, and *P2ry12*/*C1qa* was used to identify microglia. The analysis reported here focused on the glial population and layer IV neurons, for which the following numbers of cells per condition passed quality control filters and were included in subsequent analyses. For cell numbers see Supplemental Figure 9g.

### Identification of differentially-expressed genes

Differential gene expression analyses were performed in pairwise fashion between each of the four groups using the R package Monocle2^72^. The data were modeled using a negative binomial distribution consistent with data generated by high-throughput single-cell RNA-seq platforms such as inDrops. Unlike deep single-cell sequencing, inDrops probabilistically captures/samples the transcriptome of each cell and retrieves only a small fraction of all the present transcripts. Genes whose differential gene expression false discovery rate (FDR) was less than 0.10 (FDR < 0.10) were considered statistically significant.

### Cell Counts

For microglia cell counts, single plane 10x epifluorescence images were collected within the barrel cortex at the same exposure time with a Zeiss Observer microscope equipped with Zen Blue acquisition software (Zeiss; Oberkochen, Germany). Images of entire barrel fields from 10x images were quantified blind to genotype in ImageJ (NIH). Each individual barrel per field of view was outlined and grouped as a single ROI. The same ROI was transposed to the thresholded microglia channel where the number of microglia in the barrels was quantified by counting the number of cells within the total barrel ROI. The entire perimeter of the barrel field was then outlined and the number of microglia over the entire barrel field area was analyzed. The number of microglia within the septa was quantified by subtracting the total number of microglia within the barrels from the total number of microglia within the entire S1 ROI. The level of microglia infiltration into the barrel field was quantified by calculating a ratio of the total number of microglia in the barrels divided by the total number of microglia within the septa for each time point. Note, *Cx3cr1*^+/-^ littermates were used for comparison to *Cx3cr1*^-/-^ mice as microglia within both sets of mice are labeled with EGFP and show similar changes in TC synapses (Fig 2). Similar results were obtained in wild-type mice labeling microglia with Iba-1 (data not shown).

For ATF3/Caspase-3/APP positive cell counts, single plane 20x (for trigeminal nerve ganglia) and single plane 10x (for thalamic VPM and primary somatosensory cortex) were taken. The number of NeuN positive and ATF3/Caspase 3/APP positive cells for each plane were counted and divided by the total number of NeuN positive cells and normalized to the total field of view area. For peripheral macrophage infiltration to the whisker follicles following cauterization, single plane 20x images were taken. The number of CX3CR1-EGFP positive and CD45 positive cells within a given whisker follicle were counted and normalized to the total area of the whisker follicle.

### *In situ* RNA hybridization

*In situ* RNA hybridization was performed according to the manufacturer’s specification with slight modifications (ACDBio; Newark, CA). Briefly, mice were perfused with 4% PFA and brains were post-fixed for 24 hours. 10μm cryo-sections were prepared and stored in −80°C. Prior to *in situ* hybridization, cryo-sections were equilibrated to room temperature for 1 hour, then were dehydrated in a serial dilution of ethanol. Sections were incubated in hydrogen peroxide for 10 minutes and rinsed with RNase free water. Sections were treated with “Protease Plus” for 15 minutes at room temperature and rinsed with phosphate buffered saline. *In situ* probes were added and incubated for 2 hours at 40°C. Subsequent amplification steps were performed according to the manufacturer’s specification. Slices were immunostained following *in situ* hybridization. Sections were washed in 1X PBS for 10 minutes and blocked in 0.01% TritonX-100 and 2% normal goat serum for 30 minutes. Primary antibody (NeuN; Millipore) prepared in 0.01% TritonX-100 and 2% normal goat serum and slides were incubated overnight at room temperature. Secondary antibody was prepared in 0.01% TritonX-100 and 2% normal goat serum and incubated at room temperature for 2 hours. For *Cx3cl1 in situ* quantification, fluorescent RNA signal was localized to DAPI and NeuN positive or DAPI and NeuN negative cells with a MATLAB script (vR2016b). Individual channels were segmented with a set threshold for dilating the masked signal around the NeuN positive or DAPI positive channels. For *Adam10 in situ* quantification, fluorescent RNA signal was co-localized to NeuN positive or *Cx3cr1EGFP* positive cells and the number of puncta co-localized to either signal was measured with Image J (NIH).

### Structured Illumination Microscopy imaging

Immunostained sections were prepared as described above. Images were acquired on a GE Healthcare DeltaVision OMX microscope for structured illumination microscopy. Images were then processed in Imaris (Bitplane; Zurich,Switzerland) in order to 3D reconstruct the cell and visualize engulfed material.

### Slice Preparation and Electrophysiological Recordings

Male mice (∼3 months old) were anesthetized by intraperitoneal injection of sodium pentobarbital (200 mg/kg) and then were decapitated. The brain was quickly removed and placed in an oxygenated ice-cold cutting solution containing (in mM): 2.5 KCl, 1.25 NaH2PO4•H2O, 20 HEPES, 2 Thiourea, 5 Na-ascorbate, 92 NMDG, 30 NaHCO3, 25 D-Glucose, 0.5 CaCl2•2H2O and 10 MgSO4•7H2O. Brain slices (200 μM) were made using a Leica VT1200 vibratome (Leica Biosystems Inc.). The brain slices were immediately transferred into an incubation chamber containing oxygenated cutting solution at 34°C for 20 minutes. Slices were transferred into oxygenated artificial cerebrospinal fluid (ACSF) at room temperature (24°C) for recording. ACSF solution contains (in mM): 125 NaCl, 2.5 KCl, 1.2 NaH2PO4•H2O, 1.2 MgCl2•6H2O, 2.4 CaCl2•2H2O, 26 NaHCO3, 11 D-Glucose. Slices were left in this chamber for at least 1 hour before being placed in a recording chamber at room temperature. Single slices were transferred into a recording chamber continually superfused with oxygenated ACSF (30 − 32°C) at a flow rate of ∼2 ml/min for recording. Cells were visualized using infrared differential interference contrast (IR-DIC) imaging on an Olympus BX-50WI microscope. Electrophysiological recordings were recorded using an Axon Multiclamp 700B patch-clamp amplifier (Molecular Devices). Spontaneous excitatory postsynaptic current (sEPSC) was acquired in the whole cell configuration and gap-free acquisition mode in Clampex (Axon Instruments). Neurons were held at a membrane potential of −70 mV. Signals were filtered at 1 kHz using the amplifier’s four-pole, low-pass Bessel filter, digitized at 10 kHz with an Axon Digidata 1440A interface and stored on a personal computer. Pipette solution contained (in mM) 120 K gluconate; 5 KCl; 2 MgCl2•6H2O 10 HEPES; 4 ATP, 2 GTP. sEPSCs were recorded in the presence of bicuculline (20 μM) in the bath solution to block GABA_A_ receptors. After recordings stabilized, 1 min duration of recording was taken for sEPSC analysis. sEPSC frequency and amplitude was detected using Mini Analysis Program (Synaptosoft Inc. Fort Lee, NJ). All recordings and quantification were performed blind.

### Pharmacological Inhibition of ADAM10

C57Bl/6J (fluorescence intensity analysis) or Cx3cr1^EGFP/+^ (microglia engulfment analysis) mice were daily injected intraperitoneally with 25 mg/kg GI254023X (Millipore-Sigma, Darmstadt, Germany). The drug was prepared daily in 0.1M carbonate buffer/10% DMSO vehicle. Control littermate animals were vehicle injected following the same drug schedule for both fluorescence intensity and microglia engulfment analyses.

### Statistical Analysis

GraphPad Prism 7 (La Jolla, CA) provided the platform for all statistical and graphical analyses. All data sets were first tested and found to be normally distributed and parametric stats were subsequently run. Analyses included Students t-test when comparing 2 conditions or two-way ANOVA followed by Sidak’s or Tukey’s post hoc analyses (indicated in figure legends). For population data (percentage of cells expressing or binned data) a two-tailed Fisher’s exact test or a Chi-square was used (indicated in figure legends). All p and n values are specified within each figure legend.

