## Supplemental Figures for "Sensory lesioning induces microglial synapse elimination via ADAM10 and fractalkine signaling"

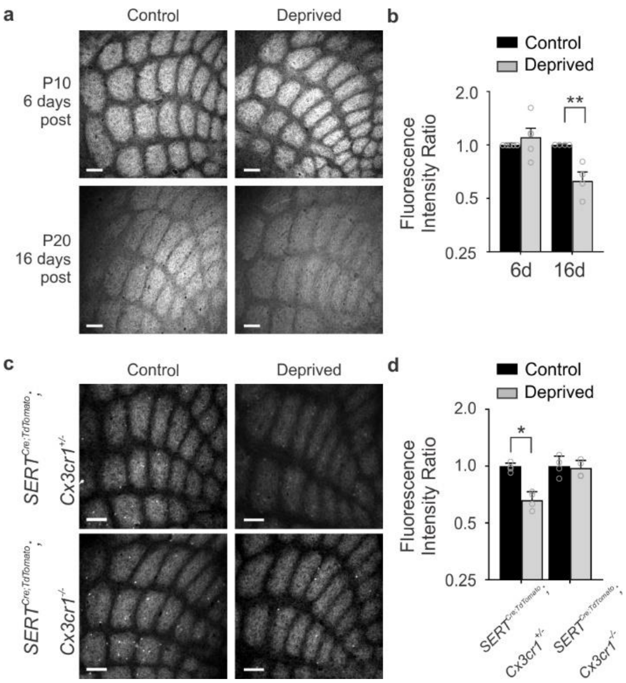


**Supplementary Figure 1. TC input elimination is observed following whisker trimming and with genetic labeling of TC inputs. a,** Daily whisker trimming from P4 results in decreased barrel fluorescence intensity by day 16 (bottom panels), but no by day 6 (top panels). Scale bar, 150 µm. **b,** Quantification for barrel fluorescence intensity in a (Two-way ANOVA with Sidak's post hoc, 6-day control vs trimmed, n = 5 animals, $P$ = 0.5093, $t$ = 1.078, $df$ = 14; 16-day control vs trimmed, n = 4 animals, $P$ = 0.0071, $t$ =3.496, $df$ = 14). Data are normalized to the control barrel cortex within the same animal for each timepoint. **c,** TC inputs labeled by transgenic expression of tdTomato are eliminated in the deprived barrel cortex in a CX3CR1-dependent manner following whisker lesioning by cauterization. Representative tangential sections of *Sert*-Cre tdTomato labeled TC inputs within layer IV of the control and deprived barrel cortices of *Cx3cr1*$^{+/-}$ (top row) and *Cx3cr1*$^{-/-}$ (bottom row) mice 7 d post-sensory deprivation. Scale bar, 150 µm. **d**, There is a significant decrease in TC inputs as measured by fluorescence intensity of tdTomato signal in the deprived (gray bars) vs. the contralateral control (black bars) barrel cortex in *Cx3cr1*$^{+/-}$ mice 7 d-post deprivation. No significant decrease in fluorescence was observed in *Cx3cr1*$^{-/-}$ mice. Data are normalized to the control barrel cortex for each genotype. (Two-Way ANOVA with Sidak's post hoc, n=4 animals per genotype, *Cx3cr1*$^{+/-}$ control vs deprived, $P$ = 0.0405, $t$ = 2.668, $df$ = 12; *Cx3cr1*$^{-/-}$ control vs deprived, $P$ = 0.9808, $t$ = 0.1782, $df$ = 12.) All data presented as mean ± SEM.

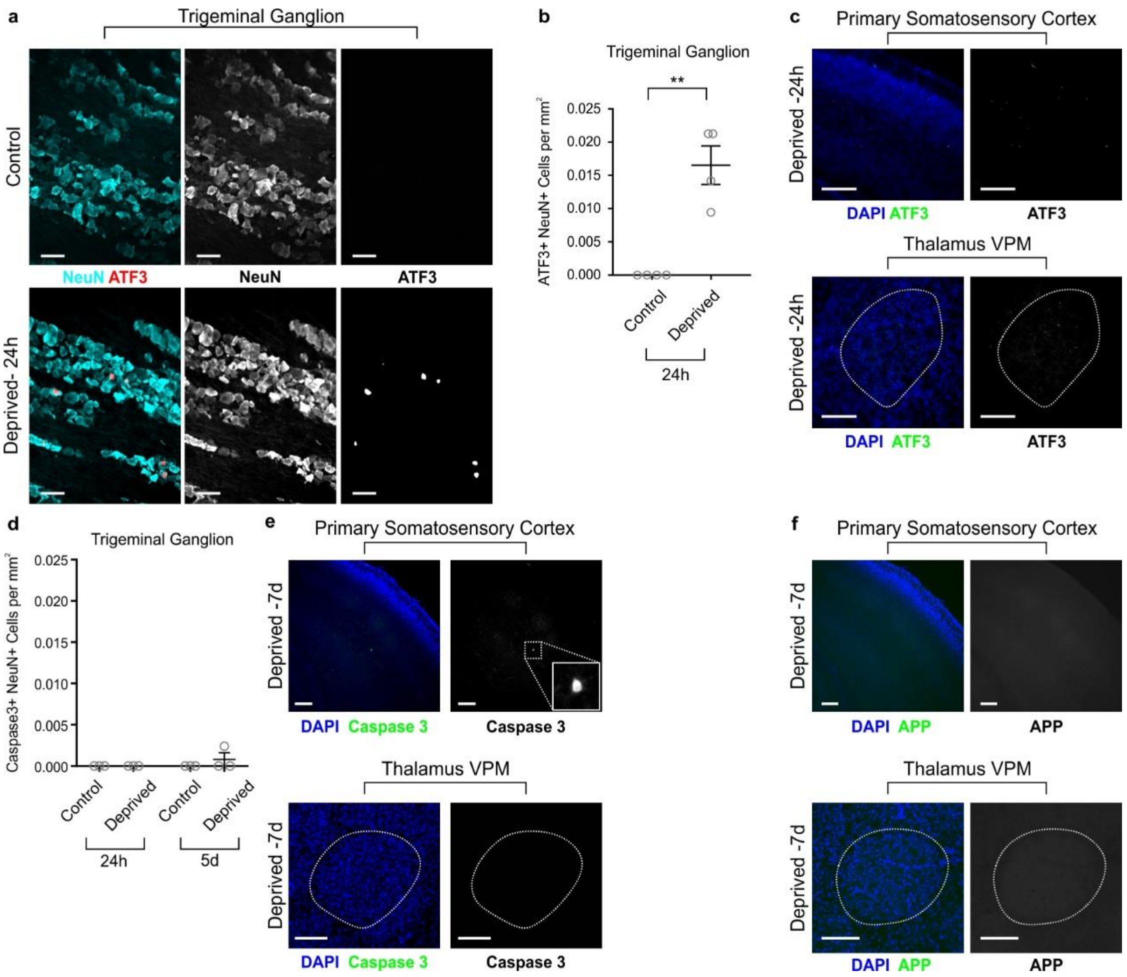


**Supplementary Figure 2. Effects of whisker lesioning at P4 on cell death, axon degeneration, and cell stress in the barrel cortex circuit. a,** Immunostaining of trigeminal ganglia, which contain the neurons that innervate the whisker follicles. There is a significant increase in ATF3 (red, marker of cell stress) in NeuN-positive neurons (cyan) at 24h post whisker lesioning (deprived, bottom row). Scale bar, 50 µm. **b**, Quantification for ATF3 signal co-localized to NeuN in the control and deprived trigeminal ganglia. (Two-tailed Student's t-Test, n = 4 animals; b, $P$ = 0.0012, $t$ = 5.715, $df$ = 6). **c**, Representative images show ATF3 signal (green) is not detected in the ventral posterior medial nucleus (VPM) of the thalamus (bottom panels; dotted line borders the VPM) nor the primary somatosensory cortex (top panels) 24 hours after whisker lesioning. Scale bars, 150 µm. **d**, Quantification for cell death marker cleaved caspase 3 shows cell death does not occur 24h nor 5d after whisker lesioning in the trigeminal ganglia. (Two-way ANOVA with Sidak's post hoc, no significant differences across comparisons; n = 3 animals per timepoint). **e-f**, Representative images for the deprived somatosensory cortex and VPM 7 days after whisker lesioning in control mice shows no increased cell death (e; caspase 3, green) nor axon degeneration (f; APP, green) in either brain region. Scale bars, 150 µm.

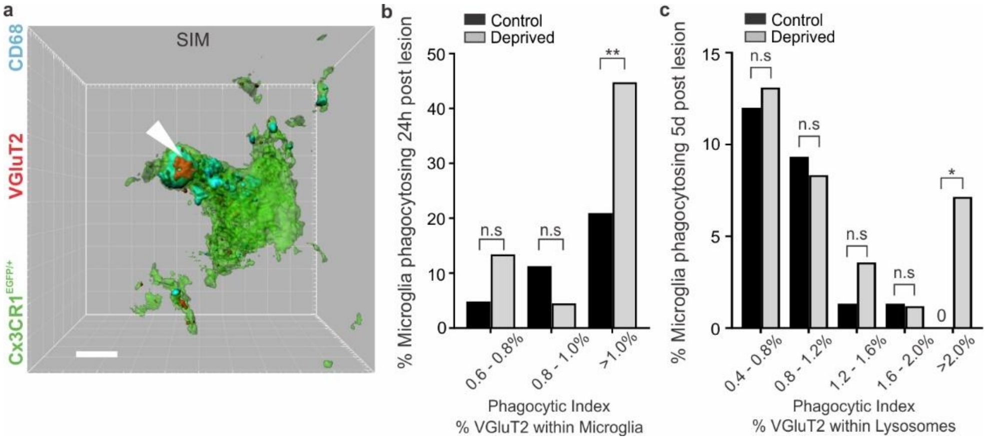


**Supplementary Figure 3. Super resolution imaging reveals TC inputs are internalized within microglia following sensory deprivation.** **a**, Structured illumination microscopy (SIM) of a microglia ($CX3CR1^{EGFP/+}$, green) which has engulfed VGluT2-positive TC presynaptic inputs (red) within its lysosomes (anti-CD68, cyan) in the deprived barrel cortex 24 h after whisker removal. Scale bar, 5 µm. **b-c**, Quantification of the % of microglia out of the total microglial population phagocytosing VGluT2-positive inputs 24h (b) and 5d (c) after whisker lesioning in the control (black bars) and deprived (grey bars) cortices. Microglia in the deprived cortex both 24h and 5d after lesioning have a higher proportion of cells with a high phagocytic index (b, >1.0%; c, >2.0%) compared to the control cortex (24h: Two-Sided Fisher's Exact Test, for >1.0% $P = 0.0051$, no significant difference for all other comparisons; 5d: Two-Sided Fisher's Exact Test, for >2.0% $P = 0.0298$, no significant difference for all other comparisons).

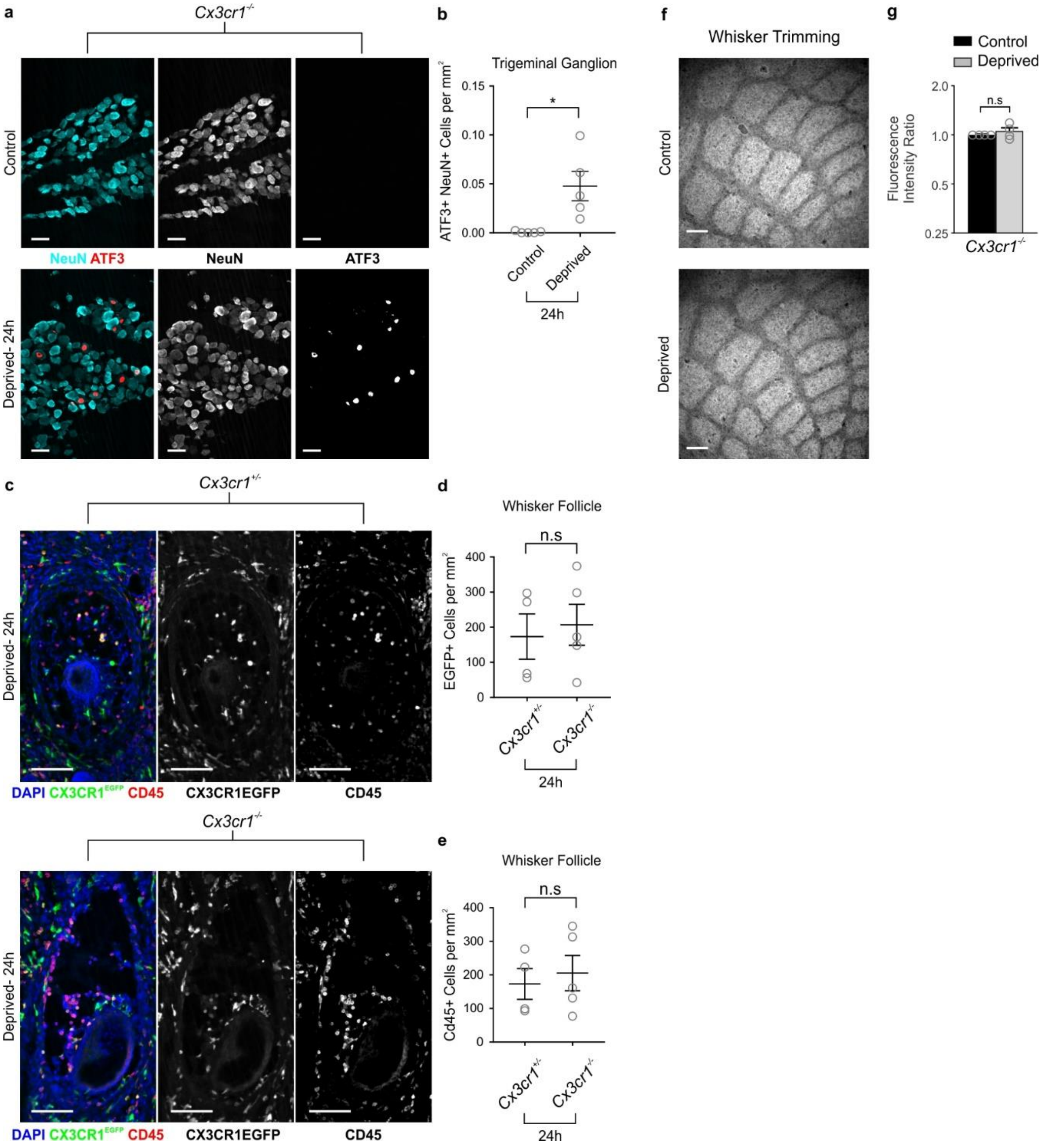


**Supplementary Figure 4. Whisker lesioning in *Cx3cr1*$^{-/-}$ animals results in similar cell stress response and wound healing at the whisker follicles compared to *Cx3cr1*$^{+/+}$ animals. a-b,** Whisker lesioning increases ATF3 signal in the ipsilateral trigeminal nerve ganglion of *Cx3cr1*$^{-/-}$ animals (deprived, bottom panels, compare to Supplementary Figure 2a). Nuclei positive for ATF3 (red) are co-localized to NeuN signal (cyan). Scale bar, 50 µm. (Two-tailed Student's t-Test, $P$ = 0.0152, $t$ = 3.362, $df$ = 6; n = 4 animals). **c-e,** Whisker lesioning in *Cx3cr1*$^{-/-}$ mice (bottom panels) results in similar wound healing response as measured by recruitment of CX3CR1-EGFP-positive (d; Two-tailed Student's t-Test, $P$ = 0.7115, $t$ = 0.3852, $df$ = 7; n = 4-5 animals) and CD45 macrophages/monocytes (e; Two-tailed Student's t-Test, $P$ = 0.6656, $t$ = 0.4511, $df$ = 7; n = 4-5 animals) compared to *Cx3cr1*$^{+/+}$ animals (top panels) 24 hours after injury. Scale bars, 100 µm. **f-g,** Whisker trimming in *Cx3cr1*$^{-/-}$ animals from P4 to P20 (16 days deprivation) does not result in a decrease in barrel fluorescence intensity (compare to Supplementary Figure 1a-b). Top panel, control barrel field fluorescence; bottom panel, deprived barrel field fluorescence at P20. Quantification for fluorescence intensity (g) does not show a significant decrease in fluorescence (Two-tailed Student's t-test, $P = 0.3455$, $t = 1.024$, $df = 6$).

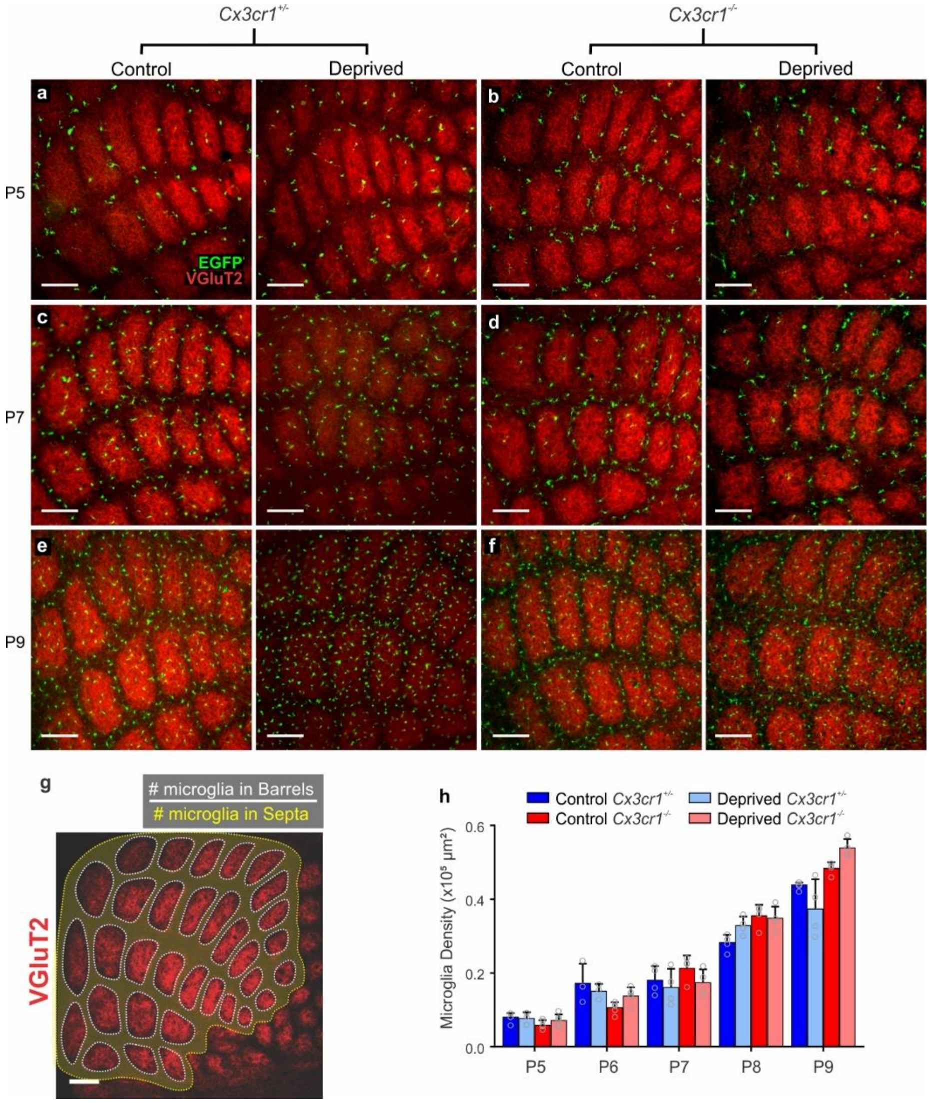


**Supplementary Figure 5. Total number of microglia across the barrel field increases with postnatal age and is similar in *Cx3cr1*$^{+/-}$ and *Cx3cr1*$^{-/-}$ mice. a-f,** Uncropped images of microglia recruitment to barrel centers from Figure 5. Scale bar, 150 µm. **g,** For each genotype, the number of microglia within the septa (yellow highlighted area) and barrels (outlined by white dotted lines) were quantified in deprived and control, non-deprived layer IV barrel cortices. A ratio was then calculated: # of microglia within the barrel divided by the # of microglia within the septa. Scale bar, 150 µm. **h**, For each postnatal age analyzed, the total microglial cell density over the entire layer IV primary somatosensory cortex is the same in *Cx3cr1*$^{+/-}$ and *Cx3cr1*$^{-/-}$ mice. (Two-Way ANOVA with Tukey's post hoc test, n=4 animals per genotype, no significant differences across comparisons). All data presented as mean ± SEM.

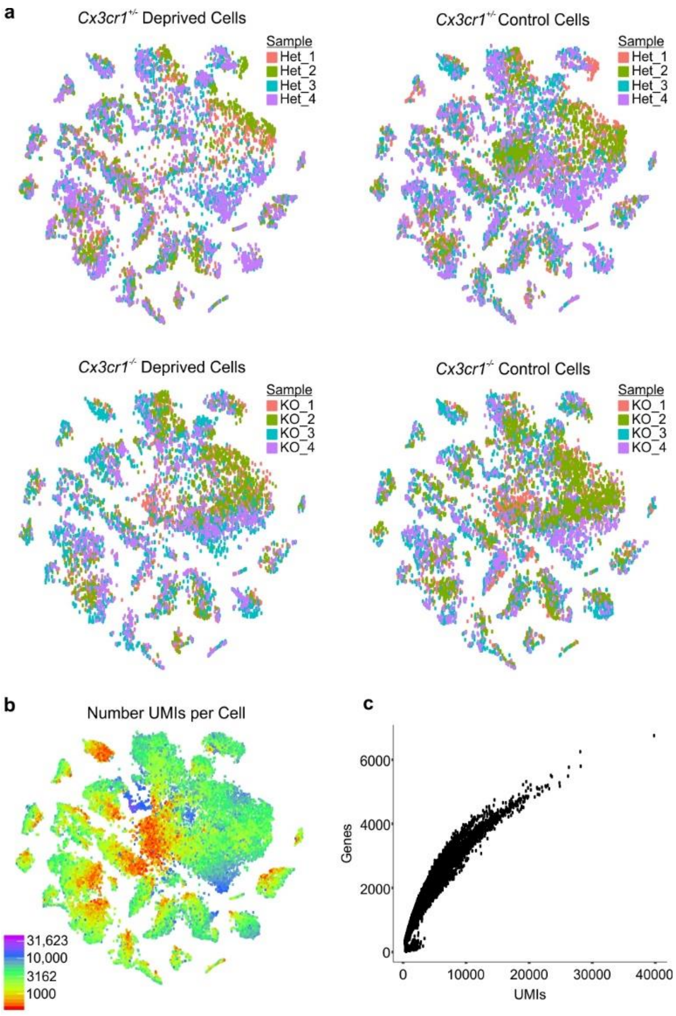


| d | Reads | Reads with unique alignment | Reads with no alignments | UMIFM |
|---|---|---|---|---|
| Mouse01 | 5131.28 | 2952.705359 | 1962.24287 | 1642.79 |
| Mouse02 | 7192.32 | 3777.569744 | 3162.182974 | 1893.78 |
| Mouse03 | 7324.4 | 4413.287984 | 2663.654332 | 1881.9 |
| Mouse04 | 7508.27 | 4705.885216 | 2555.412059 | 1605.41 |
| Mouse05 | 8392.64 | 4515.337675 | 3557.100172 | 2192.74 |
| Mouse06 | 3701.45 | 2298.715843 | 1261.619289 | 1526.6 |
| Mouse07 | 9334.52 | 5821.114634 | 3206.723902 | 2346.93 |
| Mouse08 | 5222.92 | 3202.010429 | 1843.785889 | 1687.41 |
| Mouse09 | 12628.7 | 8791.609852 | 3401.364916 | 3276.6 |
| Mouse10 | 11396.9 | 7830.870018 | 3153.854077 | 3031.85 |
| Mouse11 | 8884.22 | 6360.826473 | 2210.066183 | 2783.4 |
| Mouse12 | 14058 | 9797.345388 | 3786.499526 | 3215.88 |
| Mouse13 | 13746.8 | 9685.512426 | 3605.394877 | 3499.86 |
| Mouse14 | 11478.6 | 8073.456325 | 3011.768951 | 3179.17 |
| Mouse15 | 4042.12 | 2758.699306 | 1134.021612 | 1960.2 |
| Mouse16 | 11004.2 | 7639.631287 | 3006.15713 | 3021.2 |

**Supplementary Figure 6. Biological replicate contribution and sequencing depth of cells captured by inDrops. a,** Clustering cells from each animal condition for each biological replicate reveals even distribution of cells from each animal across all identified cell populations. Each tSNE plot represents cells from each animal genotype and condition (*Cx3cr1*$^{+/-}$ control, *Cx3cr1*$^{+/-}$ deprived, *Cx3cr1*$^{-/-}$ control, *Cx3cr1*$^{-/-}$ deprived). **b**, The number of unique molecular identifiers (UMIs) per cell processed by inDrops single-cell RNAseq (logarithmic scale). **c**, The number of genes positively identified increases with increasing UMIs. **d**, Table containing average number of reads per cell across individual animals.

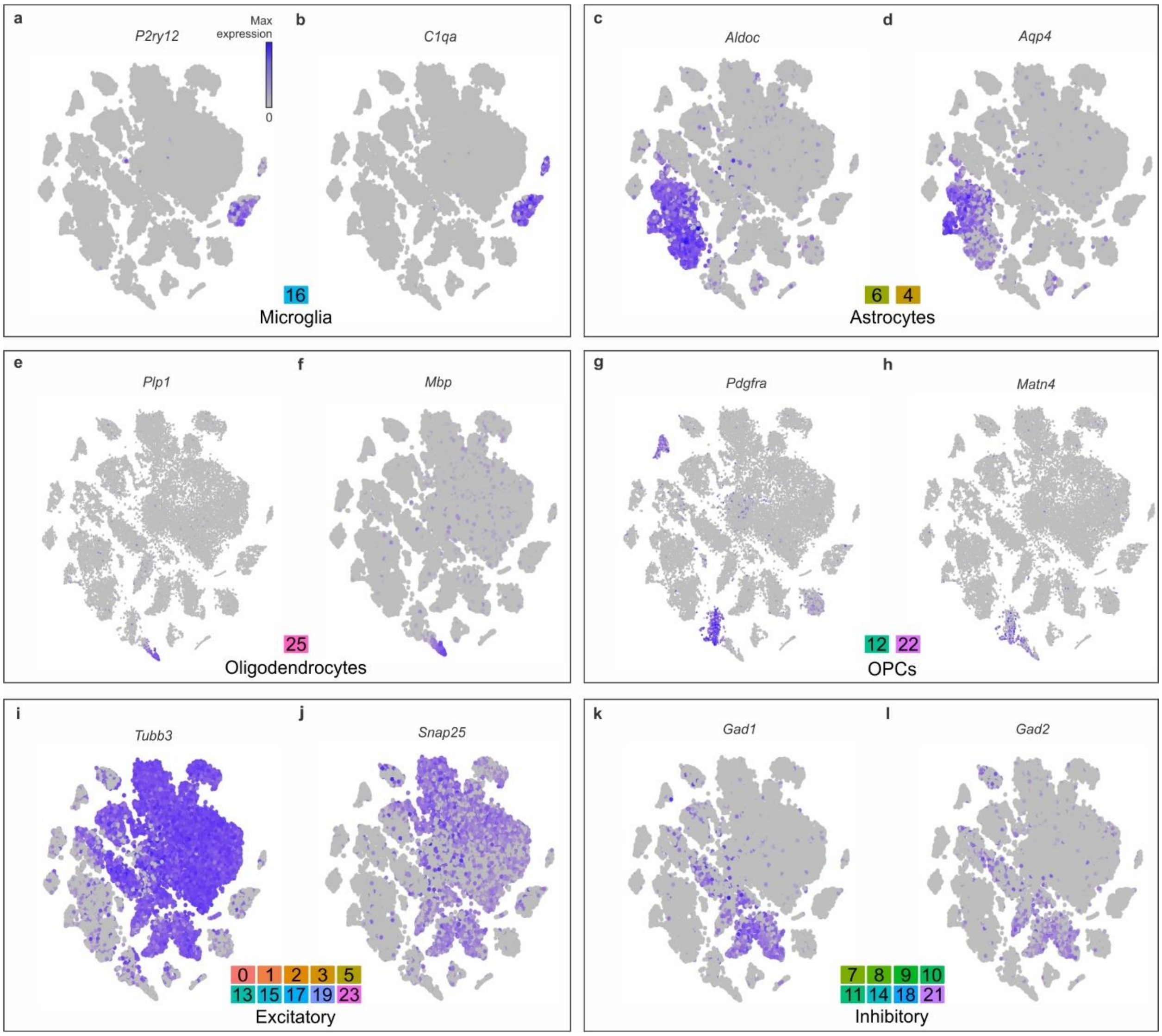


**Supplementary Figure 7. Cell populations clustered through principal component analysis were identified according to specific cell-type markers. a-b**, Microglia cluster identified by *P2ry12* and *C1qa*. **c-d**, Astrocyte clusters identified by *Aldoc* and *Aqp4* expression. **e,**f, Oligodendrocyte cluster identified by *Plp1* and *Mpb* expression. **e-f**, Oligodendrocyte precursor cell clusters identified by *Pdgfra* and *Matn4* expression. **i-j**, Neuron clusters identified by *Tubb3* and *Snap25* expression. **k-l**, Inhibitory neurons identified by *Gad1* and *Gad2* expression.

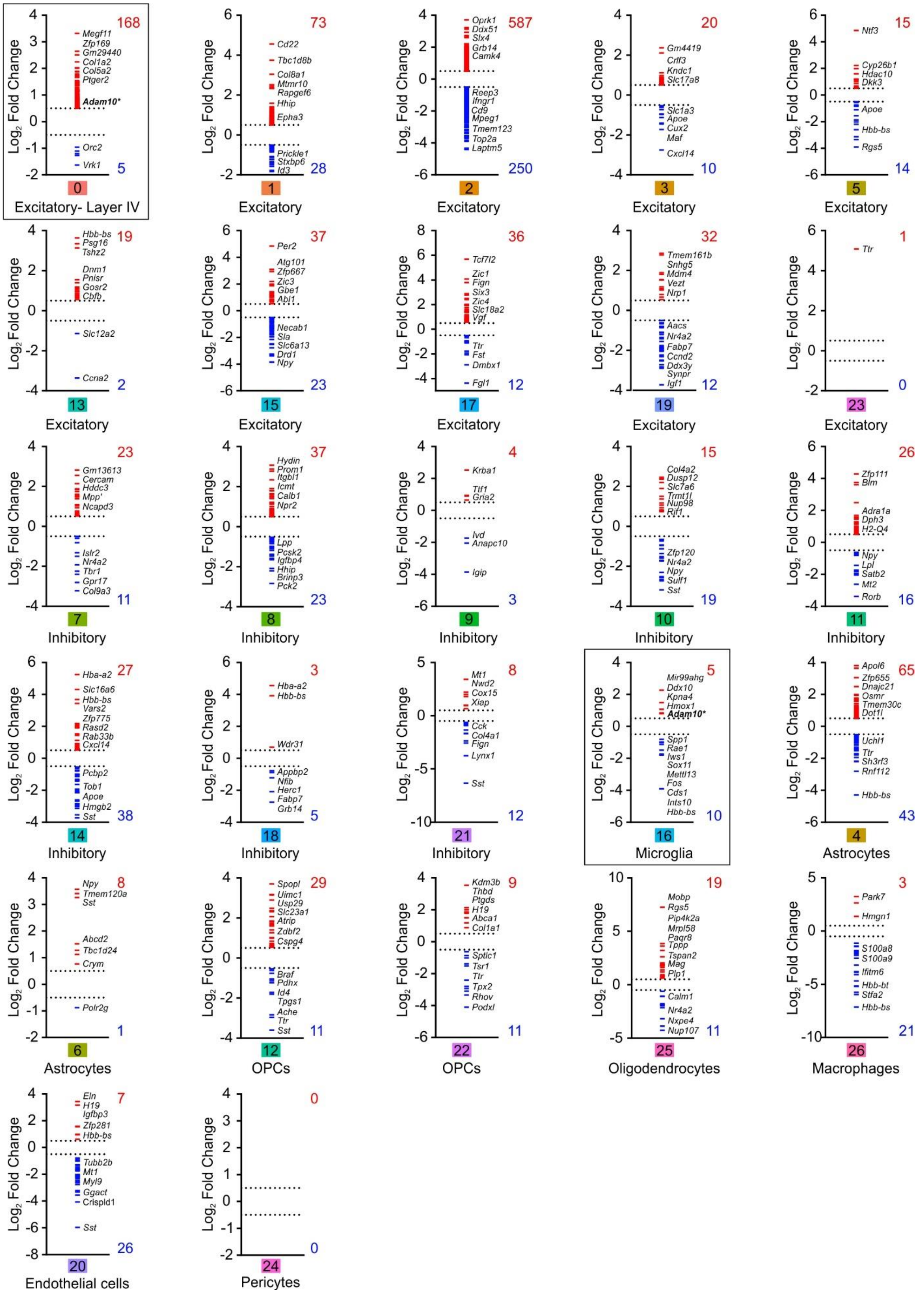
Log2 Fold Change
0 Excitatory- Layer IV
168
Megf11 Zfp169 Gm29440 Col1a2 Col5a2 Ptger2 Adam10*
Orc2 Vrk1
5
1 Excitatory
73
Cd22 Tbc1d8b Col8a1 Mtmr10 Rapgef6 Hhip Epha3
Prickle1 Stxbp6 Id3
28
2 Excitatory
587
Oprk1 Ddx51 Slx4 Grb14 Camk4
Reep3 Ifngr1 Cd9 Mpeg1 Tmem123 Top2a Laptm5
250
3 Excitatory
20
Gm4419 Crlf3 Kndc1 Slc17a8
Slc1a3 Apoe Cux2 Maf Cxcl14
10
5 Excitatory
15
Ntf3 Cyp26b1 Hdac10 Dkk3
Apoe Hbb-bs Rgs5
14
13 Excitatory
19
Hbb-bs Psg16 Tshz2 Dnm1 Pnisr Gosr2 Cbfb
Slc12a2 Ccna2
2
15 Excitatory
37
Per2 Atg101 Zfp667 Zic3 Gbe1 Abl1
Necab1 Sla Slc6a13 Drd1 Npy
23
17 Excitatory
36
Tcf7l2 Zic1 Fign Six3 Zic4 Slc18a2 Vgf
Ttr Fst Dmbx1 Fgl1
12
19 Excitatory
32
Tmem161b Snhg5 Mdm4 Vezt Nrp1
Aacs Nr4a2 Fabp7 Ccnd2 Ddx3y Synpr Igf1
12
23 Excitatory
1
Ttr
0
7 Inhibitory
23
Gm13613 Cercam Hddc3 Mpp7 Ncapd3
Islr2 Nr4a2 Tbr1 Gpr17 Col9a3
11
8 Inhibitory
37
Hydin Prom1 Itgbl1 Icmt Calb1 Npr2
Lpp Pcsk2 Igfbp4 Hhip Brinp3 Pck2
23
9 Inhibitory
4
Krba1 Ttf1 Gria2
Ivd Anapc10 Igip
3
10 Inhibitory
15
Col4a2 Dusp12 Slc7a6 Trmt1l Nup98 Rif1
Zfp120 Nr4a2 Npy Sulf1 Sst
19
11 Inhibitory
26
Zfp111 Blm Adra1a Dph3 H2-Q4
Npy Lpl Satb2 Mt2 Rorb
16
14 Inhibitory
27
Hba-a2 Slc16a6 Hbb-bs Vars2 Zfp775 Rasd2 Rab33b Cxcl14
Pcbp2 Tob1 Apoe Hmgb2 Sst
38
18 Inhibitory
3
Hba-a2 Hbb-bs Wdr31
Appbp2 Nfib Herc1 Fabp7 Grb14
5
21 Inhibitory
8
Mt1 Nwd2 Cox15 Xiap
Cck Col4a1 Fign Lynx1 Sst
12
16 Microglia
5
Mir99ahg Ddx10 Kpna4 Hmox1 Adam10*
Spp1 Rae1 Iws1 Sox11 Mettl13 Fos Cds1 Ints10 Hbb-bs
10
4 Astrocytes
65
Apol6 Zfp655 Dnajc21 Osmr Tmem30c Dot1l
Uchl1 Ttr Sh3rf3 Rnf112 Hbb-bs
43
6 Astrocytes
8
Npy Tmem120a Sst Abcd2 Tbc1d24 Crym
Polr2g
1
12 OPCs
29
Spopl Uimc1 Usp29 Slc23a1 Atrip Zdbf2 Cspg4
Braf Pdhx Id4 Tpgs1 Ache Ttr Sst
11
22 OPCs
9
Kdm3b Thbd Ptgds H19 Abca1 Col1a1
Sptlc1 Tsr1 Ttr Tpx2 Rhov Podxl
11
25 Oligodendrocytes
19
Mobp Rgs5 Pip4k2a Mrpl58 Paqr8 Tppp Tspan2 Mag Plp1
Calm1 Nr4a2 Nxpe4 Nup107
11
26 Macrophages
3
Park7 Hmgn1
S100a8 S100a9 Ifitm6 Hbb-bt Stfa2 Hbb-bs
21
20 Endothelial cells
7
Eln H19 Igfbp3 Zfp281 Hbb-bs
Tubb2b Mt1 Myl9 Ggact Crispld1 Sst
26
24 Pericytes
0
0

**Supplementary Figure 8. Genes significantly changed for each cell population in the deprived cortex of *Cx3cr1*$^{+/-}$ animals.** Genes with significant changes in expression (FDR <0.10) and $Log_2$ Fold Change greater than 0.5 or less than -0.5 in the deprived cortex are plotted for every identified cell population. Note ADAM10 (bold text and asterisks) is only significantly increased in layer IV excitatory neurons and microglia (outlined graphs).

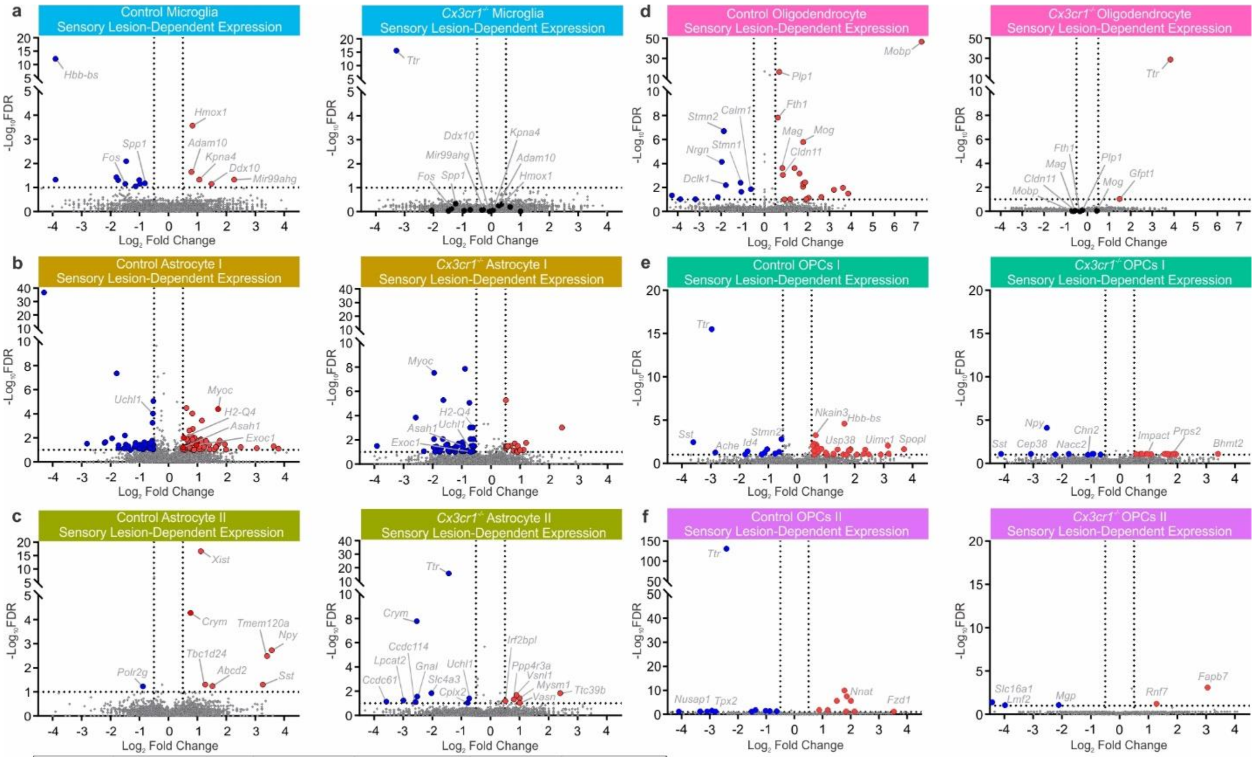


**g**

| Cluster # | Cell type | HET Control | HET Deprived | KO Control | KO Deprived |
|---|---|---|---|---|---|
| 0 | Excitatory | 783 | 549 | 847 | 803 |
| 1 | Excitatory | 773 | 459 | 946 | 804 |
| 2 | Excitatory | 1135 | 415 | 873 | 438 |
| 3 | Excitatory | 697 | 562 | 781 | 724 |
| 4 | Astrocytes | 668 | 671 | 554 | 519 |
| 5 | Excitatory | 393 | 245 | 563 | 448 |
| 6 | Astrocytes | 445 | 474 | 348 | 346 |
| 7 | Inhibitory | 382 | 324 | 465 | 343 |
| 8 | Inhibitory | 411 | 392 | 329 | 362 |
| 9 | Inhibitory | 490 | 476 | 269 | 242 |
| 10 | Inhibitory | 422 | 368 | 309 | 275 |
| 11 | Inhibitory | 343 | 281 | 313 | 232 |
| 12 | OPCs | 290 | 311 | 209 | 197 |
| 13 | Excitatory | 258 | 241 | 209 | 230 |
| 14 | Inhibitory | 235 | 260 | 270 | 123 |
| 15 | Excitatory | 165 | 184 | 292 | 234 |
| 16 | Microglia | 264 | 282 | 166 | 116 |
| 17 | Excitatory | 180 | 287 | 195 | 37 |
| 18 | Inhibitory | 188 | 180 | 188 | 99 |
| 19 | Excitatory | 109 | 180 | 161 | 203 |
| 20 | Endothelial Cells | 109 | 182 | 145 | 117 |
| 21 | Inhibitory | 87 | 112 | 211 | 84 |
| 22 | OPCs | 89 | 169 | 65 | 73 |
| 23 | Excitatory | 80 | 83 | 134 | 99 |
| 24 | Pericytes | 116 | 153 | 55 | 55 |
| 25 | Oligodendrocytes | 96 | 109 | 58 | 47 |
| 26 | Macrophages | 64 | 78 | 53 | 33 |
| 27 | Unknown | 62 | 60 | 23 | 59 |
| | SUM | 9334 | 8087 | 9031 | 7342 |

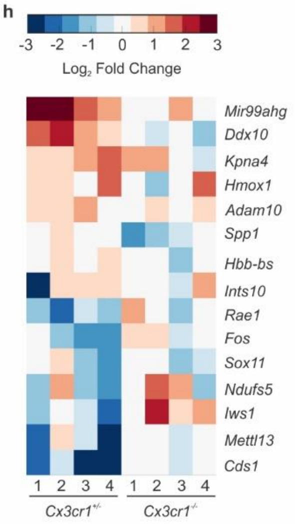


**Supplementary Figure 9. Sensory-lesion dependent gene expression changes in all major glial cell populations in the deprived cortices of *Cx3cr1*$^{+/-}$ and *Cx3cr1*$^{-/-}$ animals. a,** Gene expression changes for *Cx3cr1*$^{+/-}$ microglia (left panel) and *Cx3cr1*$^{-/-}$ microglia (left panel). Dotted lines indicates –$\log_{10}$FDR <0.10 and $\text{Log}_2$ Fold Change > 0.5 and < -0.5. **b-f,** Gene expression changes for all other identified glial cell types. **g,** Table containing the number of cells sequenced for each cell type for each animal genotype and condition. **H**, Heatmap for the

$Log_2$Fold Change for microglial genes changed across *Cx3cr1*$^{+/-}$ and *Cx3cr1*$^{-/-}$ biological replicates**.**

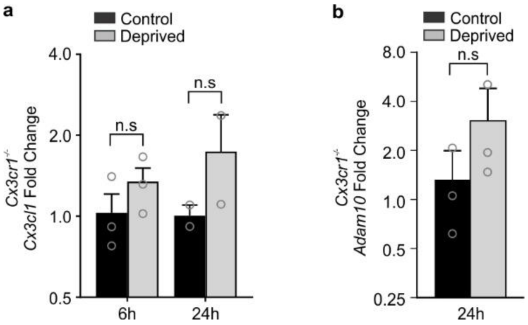


**Supplementary Figure 10. qPCR in *Cx3cr1*$^{-/-}$ primary somatosensory cortex reveals no significant increase in *Cx3cl1* nor *Adam10* expression after whisker lesioning. a**, Quantification for *Cx3cl1* expression 6h and 24h post lesioning. (Two-way ANOVA with Sidak's post hoc, no significant differences across comparisons; n = 3 animals per timepoint). **b**, Quantification for *Adam10* expression 24h post lesioning. (Two-tailed Student's t-Test, *P* = 0.4414, *t* =  0.9099, *df* = 4; n = 3 animals).

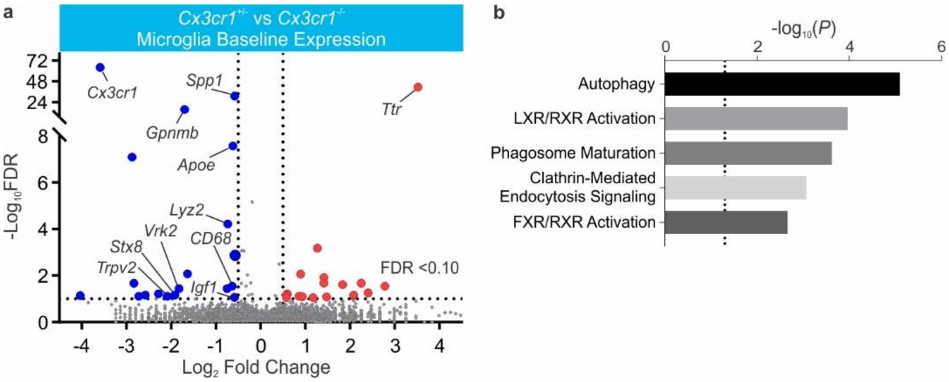


**Supplementary Figure 11. Baseline expression in *Cx3cr1*$^{-/-}$ microglia reveals significant changes in genes related to phagocytic signaling. a**, Volcano plot of genes significantly downregulated (fold change < -0.5, FDR <0.10, p<0.0005) in *Cx3cr1*$^{-/-}$ microglia within the non-deprived, control barrel cortex compared to *Cx3cr1*$^{+/-}$ littermates reveals several genes are dysregulated basally in *Cx3cr1*$^{-/-}$ microglia. **b**, Gene ontology clustering of genes in *Cx3cr1*$^{-/-}$ microglia (Dotted line, p<0.05) reveals several genes related to phagocytic signaling are significantly changed. Data analyzed through the use of IPA (QIAGEN Inc., https://www.qiagenbioinformatics.com/products/ingenuitypathway-analysis).
